# Intertwined Autophagy and Integrin Dynamics Shape Axon Growth and Regeneration

**DOI:** 10.64898/2026.08.04.742728

**Authors:** Anda Cimpean, Jessica C.F. Kwok, James W. Fawcett, Pavla Jendelová

## Abstract

Autophagy is a central pathway for cellular homeostasis, mediating degradation and recycling of cytoplasmic components through lysosomal processing. Although autophagy has been implicated in axon growth and neuronal injury responses, its role in axon regeneration remains incompletely understood. In parallel, integrin trafficking and focal adhesion dynamics are critical determinants of axonal growth, and increasing evidence indicates that autophagy regulates focal adhesion turnover in non-neuronal cells; however, whether this mechanism operates in neurons remains unknown. Here, we investigated the dynamics of autophagy during axon growth and regeneration in adult sensory neurons and examined its relationship with integrin trafficking. Using live imaging, we analyzed autophagic vesicles and integrin-containing compartments in axons under basal conditions and following axotomy. We find that axonal injury induces marked short- and long-term alterations in distal axon and growth cone autophagic vesicle dynamics, integrin trafficking, and autophagy-associated integrin turnover. Importantly, these changes correlate with axonal growth and regenerative capacity. Furthermore, we identify a functional interplay between autophagy and integrins, suggesting bidirectional regulation in which autophagy contributes to adhesion receptor recycling, while integrins feedback to modulate autophagy. Finally, pharmacological modulation indicates that autophagy plays a critical role in both axonal growth and regeneration, and that rapamycin enhances regenerative responses, likely through modulation of autophagy-dependent integrin recycling rather than a general increase in autophagic activity. Together, these findings reveal a bidirectional feedback system between integrin-mediated extracellular cues and autophagy-dependent intracellular trafficking that jointly orchestrates axon growth and regeneration.

## INTRODUCTION

One of the major challenges in neuroscience is the regeneration of adult damaged axons. Axon regeneration is complex and tightly regulated by both extrinsic cues and intrinsic neuronal properties, which together drive extensive changes in neuronal homeostasis [1]. A central regulator of neuronal homeostasis is the evolutionarily conserved autophagy pathway. Macroautophagy (hereafter autophagy) involves the engulfment of intracellular material, either in bulk or selectively via cargo receptors, into double-membrane vesicles termed autophagosomes (APs). These mature through fusion with lysosomes to form autolysosomes (ALs), where cargo is degraded and recycled to support metabolic and anabolic demands [2]. Thus, autophagy functions as a quality-control and recycling system that preserves cellular integrity while also sustaining cellular metabolism [3]. Neurons are particularly dependent on autophagy due to their post-mitotic state, extreme longevity, polarity, and high energetic requirements [4].In these cells, AP are generated predominantly in distal axons and undergo retrograde transport toward the soma while progressively maturing. This spatial organization highlights the importance of vesicular trafficking and compartmentalized regulation of autophagy in neurons [5–7]. However, despite its essential role in neuronal homeostasis, the function of autophagy in axon regeneration remains incompletely understood. Both enhancement and inhibition of autophagy have been reported to either promote or impair axonal regeneration depending on the experimental context, and impairments in AP maturation inhibits regeneration [8]. Current literature establishes a role for autophagy in axon growth and regeneration, but there is a complex and context-dependent relationship that remains unresolved [9,10].

A key process potentially linking autophagy to axon regeneration is the dynamic regulation of adhesion and membrane recycling, a connection that remains largely unexplored. Axonal regeneration is strongly associated with integrin expression and turnover [11,12]. Integrins are α/β heterodimeric receptors that cluster at the plasma membrane to form focal adhesions (FA), multiprotein complexes that couple the extracellular matrix to the actin cytoskeleton and coordinate adhesion, migration, and cytoskeletal remodeling. Distinct neuronal populations display specific integrin repertoires, with diversity largely driven by differential expression of α subunits pairing with β1 integrins. Importantly, integrin localization is tightly controlled in both space and time through endosomal trafficking pathways regulated by Rab GTPases [13,14]. Accordingly, manipulation of integrin expression or trafficking has emerged as a strategy to enhance axonal regeneration [14–17]. Recent evidence has shown that autophagy can selectively target FA components for degradation, a process termed “focal adhesion autophagy” (FA-phagy) [18]. FA-phagy has been implicated in promoting cell migration in multiple contexts [19]. Although axon regeneration and cell migration are distinct biological processes, they share fundamental mechanistic features, as the regenerating axon functions as a highly motile structure that relies on coordinated cytoskeletal remodeling, membrane trafficking, and adhesion dynamics to extend toward its target [20]. Given these parallels, it is plausible that autophagy similarly contributes to axonal growth and regeneration by regulating integrin availability and adhesion turnover in neurons.

In this study, we aimed to characterize autophagy dynamics during axon growth and regeneration and to determine whether autophagy regulates integrin recycling at the distal axon and growth cone, thereby influencing axonal extension. Using live imaging of adult sensory neurons, we analyze the dynamics of autophagic and integrin-containing vesicles. Our quantitative analyses reveal that vesicle dynamics correlate with axonal growth and regenerative capacity. Furthermore, we show that axotomy induces both acute and long-term changes in axonal intracellular trafficking. We identify autophagy as a regulator of integrin recycling and reveal a reciprocal interplay between these pathways. Finally, pharmacological modulation of autophagy indicates that rapamycin enhances axonal regenerative capacity, likely through modulation of autophagy-dependent integrin recycling rather than a general increase in autophagic activity.

## RESULTS

Statistics for the results section are shown in supplementary tables 1-8. In the first part of the study, we observed the location and movement of AP/AL and integrins in the growth cones and distal axons of adult DRG neurons *in vitro*. We then investigated their behavior in axotomized axons, asking whether their dynamics correlated with successful or failed axon regeneration. We used adult C57BL/6-Tg(CAG-RFP/EGFP/Map1lc3b)1Hill/J mice expressing a tandem fluorescent LC3B reporter, enabling visualization of AP and AL based on pH-dependent fluorescence [21,22]. Consequently, APs display both EGFP and RFP fluorescence (appearing yellow), while ALs exhibit only RFP signal due to EGFP quenching.

To simultaneously assess β1 integrin distribution, a far-red–conjugated antibody was used. Imaging for 2 minutes at 1 second intervals provided good data on vesicle movement. Because simultaneous three-channel imaging induced phototoxicity, experiments were performed using either EGFP-RFP or RFP-far-red configurations.

### 1. Spatial organization and dynamics of autophagic and integrin vesicles in the growth cone

Growth cones have distinct organizational and trafficking properties, with central and peripheral domains separated by a transition zone [23].

We observed that LC3⁺ autophagic vesicles (including both AP and AL) are predominantly localized within the central domain of the growth cone. However, AL were also detected in the peripheral domain and within filopodia, extending toward the leading edge of membrane protrusions. In these regions, yellow APs were frequently seen maturing into red-only ALs (Fig. 1A). Autophagic vesicles exhibited discontinuous trajectories, characterized by abrupt pauses and frequent directional changes, likely reflecting transient interactions with the cytoskeleton, other organelles, or local structural constraints (Video S1). There was continuous movement of LC3⁺ vesicles into and out of the growth cones into the axons.

**Figure 1.**
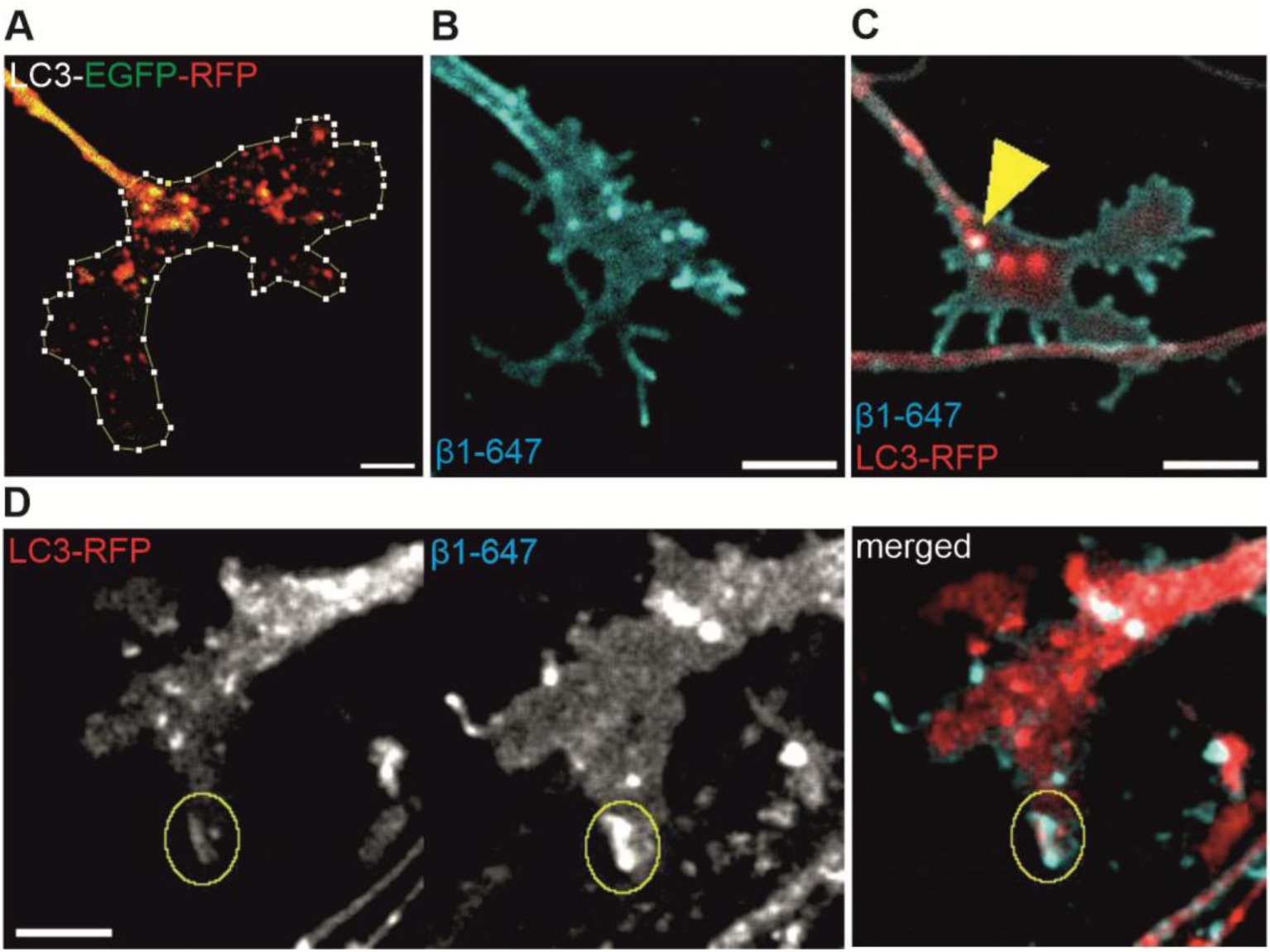
Autophagic and integrins dynamics in adult DRG growth cones. (**A**) APs (GFP+-RFP+) are predominantly localized to the central domain, whereas ALs (RFP+ only) extend into the peripheral domain, including filopodia. (**B**) β1 integrin distribution within the growth cone is primarily localized to patches at the plasma membrane. (**C**) Colocalization between β1 integrins and autophagic vesicles is largely restricted to the central domain. (**D**) Waves of LC3 and β1 integrin signals colocalize at sites of plasma membrane remodeling. Scale bar, 5 µm.

β1 integrin appeared as discrete patches at the growth cone surface and was also associated with intracellular vesicles (Fig. 1B). These structures could remain stationary at the membrane, undergo lateral diffusion, or traffic to and from more distal membrane regions (Video S2).

Simultaneous visualization of LC3⁺ and β1⁺ vesicles revealed that integrins were mostly associated with vesicles that were not LC3⁺. Those are probably recycling endosomes that mediate the majority of integrin trafficking [14]. Although LC3⁺ and β1⁺ vesicles frequently exhibited close spatial proximity and coordinated movement, true LC3-β1-integrin colocalization was rare and confined to the central domain of the growth cone, where vesicles appeared to enter or exit the axon (Fig. 1C; Video S3).

However, we more frequently observed events in which β1-integrin appeared to be removed from the plasma membrane by LC3⁺ structures. LC3 and β1-integrin occasionally organized into dynamic, wave-like structures, transiently enriching and colocalizing at sites of plasma membrane remodeling. As these waves propagated, a subset of integrin appeared to be incorporated into vesicular structures, suggesting that LC3⁺ compartments may participate in integrin recycling from the plasma membrane (Fig. 1D; Video S4). Similar LC3-associated membrane remodeling events have previously been described in migrating cancer cells [24,25]. In summary, autophagic vesicles are primarily concentrated in the central domain of the growth cone but are also present in the periphery, with AL trajectories extending into filopodia. β1 integrin is mainly localized as patches at the plasma membrane. Colocalization between β1 integrin and LC3 is limited and largely confined to vesicular structures in the central domain, whereas transient wave-like structures emerge at sites of membrane remodeling and integrin internalization.

### 2. Autophagic and integrin vesicles behavior in axon growth

To examine autophagic and integrin vesicle dynamics during physiological axon growth, imaging was focused on the distal axon segment, approximately 100 µm from the axon tip. Two imaging sessions were performed 2 hours apart, allowing axons to be classified as either growing or static (Fig. 2A; Video S5–S6).

**Figure 2.**
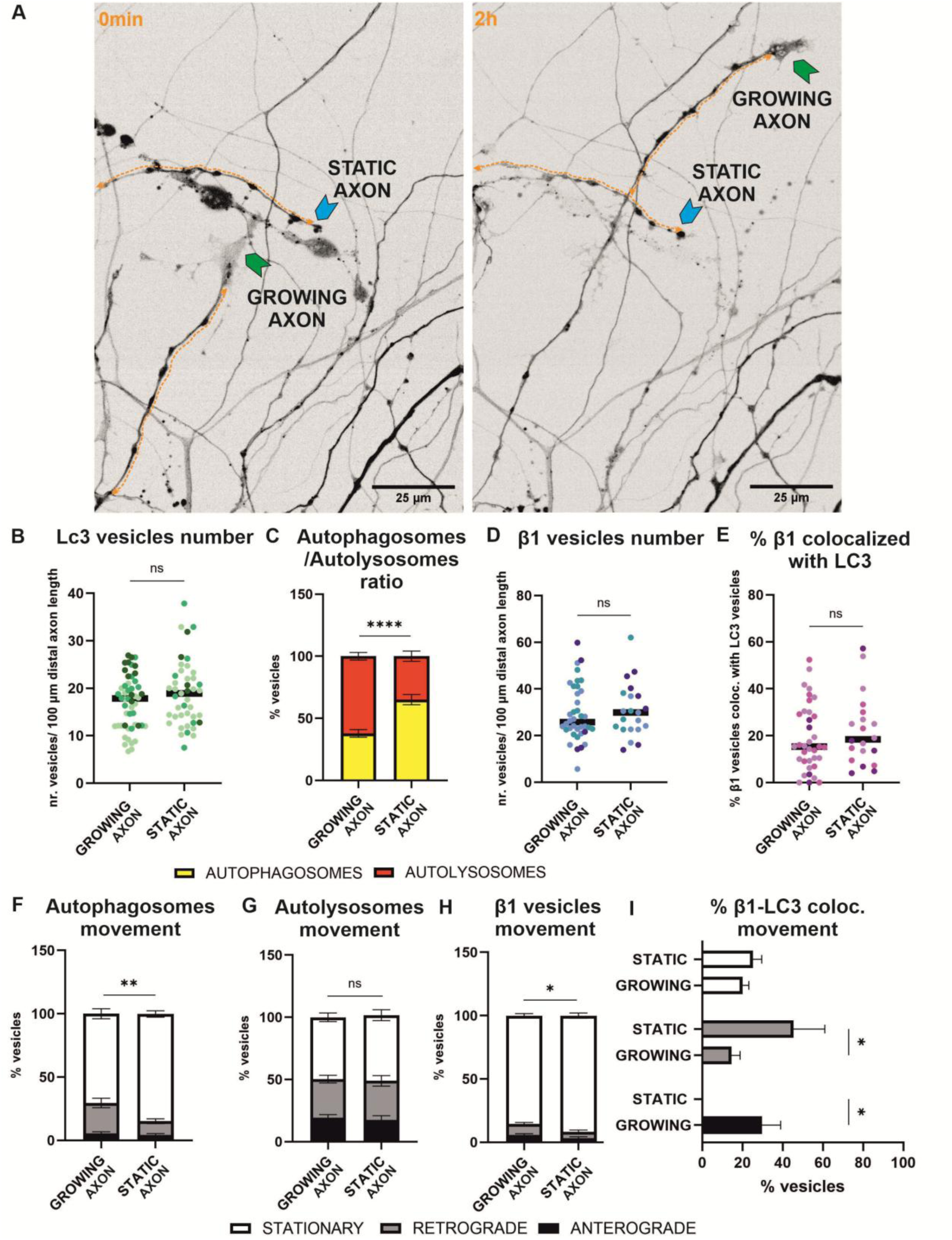
Autophagic and integrins dynamics in growing and static axons. (**A**) Representative single-channel (RFP) images from live imaging acquired 2 h apart under physiological conditions. Green arrows indicate growing axons; blue arrows indicate static axons. Orange double arrows indicate the axonal distal segment analyzed. (**B**) Total number of LC3^+^ vesicles. (**C**) Proportion of AP:AL within the total autophagic vesicle pool. (**D**) Total number of β1^+^ vesicles. (**E**) Percentage of β1^+^ vesicles colocalizing with LC3^+^ vesicles. (F–I) Trafficking analysis showing the percentage of vesicles moving anterogradely, retrogradely, or remaining stationary for APs (**F**), ALs (**G**), β1 vesicles (**H**), and β1^+^-LC3^+^ vesicles (**I**). Each dot represents a single axon. Colors indicate independent biological replicates (different animals and imaging days). 3 independent experiments were performed for each configuration (red+green configuration, n = 50 growing axons, n = 43 static axons; red+far-red configuration, n = 38 growing axons, n = 21 static axons). Bars represent mean ± SEM. Statistical analysis was performed using unpaired t-tests (B, D, E) or two-way ANOVA (C, F–I). *p < 0.05, **p < 0.01, ****p < 0.0001.

Vesicle dynamics differed significantly between these groups, but the overall abundance of vesicles did not. The total number of LC3⁺ autophagic vesicles (APs and ALs) per distal 100 µm did not differ between growing and static axons (Fig. 2B), nor did the number of β1⁺ vesicles (Fig. 2D). Along the distal axon, colocalization between LC3⁺ and β1⁺ vesicles was more frequent than in growth cones, with approximately 20% of β1⁺ vesicles associating with LC3⁺ vesicles in both conditions (Fig. 2E). In contrast, AP maturation differed substantially between the two axonal states. Growing axons exhibited a predominance of ALs relative to APs, whereas static axons showed the opposite distribution (Fig. 2C).

Vesicle trajectories were classified as anterograde (>5 µm/2 min toward the axon tip), retrograde (>5 µm/2 min toward the soma), or stationary (<5 µm/2 min). In growing axons, APs were predominantly stationary (70%), with smaller fractions undergoing retrograde (25%) or anterograde (5%) transport. In static axons, the stationary fraction was higher (85%), while retrograde transport was lower (11%), with no significant difference in anterograde movement (4%) (Fig. 2F). In contrast, ALs were more dynamic than APs and their movement did not differ between the two groups (Fig. 2G). β1⁺ vesicles were largely stationary during the 2 min live imaging period in both conditions; however, static axons showed a modest but significantly higher percentage in the stationary fraction (Fig. 2H). We next analyzed the dynamics of LC3⁺-β1⁺ colocalized vesicles. In growing axons, approximately 30% of anterogradely moving, 15% of retrogradely moving, and 20% of stationary β1⁺ vesicles were colocalizing with LC3⁺ vesicles. In static axons, colocalization was significantly enriched in the retrograde direction and absent in the anterograde direction, while the stationary fraction remained unchanged (Fig. 2I). Notably, despite similar overall levels of colocalization, and although both AP and β1 integrin vesicles exhibit reduced retrograde transport in static axons, their colocalization becomes preferentially associated with retrograde movement. This suggests a selective change in autophagic cargo trafficking rather than a global alteration in vesicle direction. Statistical details are provided in Table S1.

In summary, vesicle dynamics correlate with axonal state. Static axons differ from growing axons by a reduced number of ALs and decreased retrograde transport of APs, suggesting lower maturation. β1 integrin vesicles also show reduced retrograde transport. Autophagic involvement in integrin recycling appears more prominent in the distal axon than in the growth cone, with ∼20% of integrins trafficked/recycled via autophagic vesicles. Static axons present a redistribution of integrin–autophagy colocalization toward retrograde trafficking, with absence of anterograde trafficking. Collectively, these results show that trafficking behavior differs depending on axonal growth competence.

### 3. Axotomy alters autophagic and integrin vesicles dynamics

After characterizing vesicle behavior in physiological conditions, we sought to determine the immediate and later effects of axotomy. To explore this, we performed live imaging microscopy immediately following axotomy and then again two hours later (Video S7-8). The two-hour time point was chosen based on previous findings indicating that adult sensory axons will be regenerating by this time if they are going to [26]. This enabled retrospective classification of injured axons as regenerating (elongating by ≥5 µm within 2 h), static (axonal length change <5 µm within 2 h), or retracting (shortening by ≥5 µm within 2 h). Although all axons initially underwent a similar retraction response and formed a retraction bulb to seal the membrane, this classification allowed us to determine whether early changes in vesicle dynamics predicted the eventual regenerative outcome (Fig. 3A).

**Figure 3.**
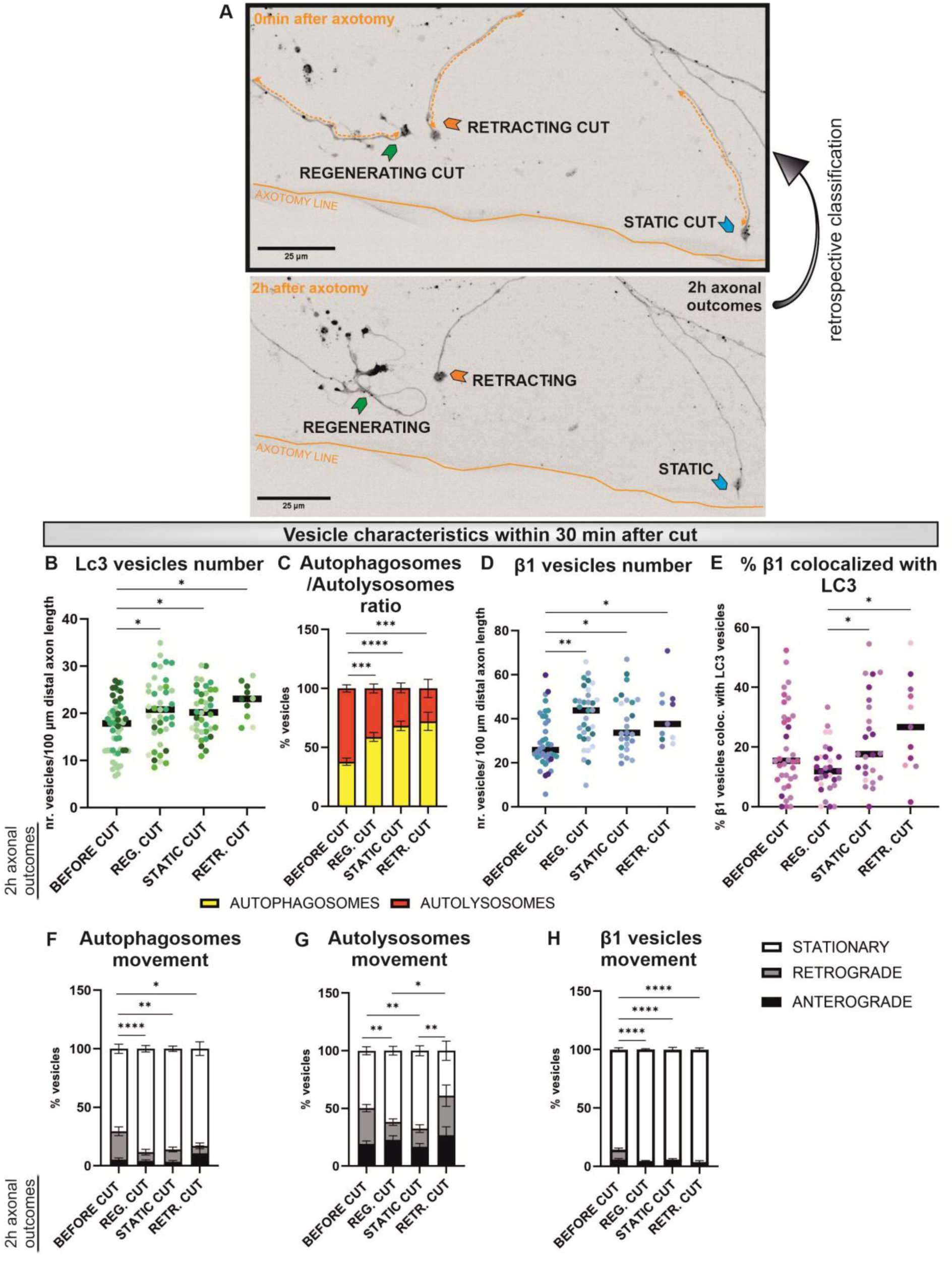
Autophagic and integrins dynamics immediately after injury. (**A**) Representative single-channel (RFP) images from live imaging acquired 2 h apart following axotomy. Axons were retrospectively classified as regenerating, retracting, or static. Green arrows indicate regenerating axons; blue arrows indicate static axons; orange arrows indicate retracting axons. Orange double arrows indicate the axonal distal segment analyzed. (**B–H**) Quantifications as in Fig. 2. Each dot represents a single axon. Colors indicate independent biological replicates (different animals and imaging days). 8 independent experiments were performed (red+green configuration, n = 41 regenerating axons, n = 38 static axons, n = 10 retracting axons; red+far-red configuration, n = 33 regenerating axons, n = 25 static axons, n = 11 retracting axons). Bars represent mean ± SEM. Statistical analysis was performed using one-way ANOVA (B, D, E) or two-way ANOVA (C, F–H). *p < 0.05, **p < 0.01, ***p < 0.001, ****p < 0.0001.

Axotomy led to an immediate increase in the total number of LC3⁺ vesicles in all distal axons, regardless of their eventual outcome (Fig. 3B). In parallel, all injured axons exhibited an immediate shift in the AP:AL ratio, with an increased proportion of APs (Fig. 3C). Similarly, β1⁺ vesicles increased in all injured axons compared to pre-injury levels (Fig. 3D). Although LC3⁺-β1⁺ colocalized vesicles did not show a significant overall increase compared to pre-injury levels, higher levels of colocalization were observed in axons that will remain static or will retract compared to those that will regenerate (Fig. 3E).

Vesicle movement changed immediately following axotomy with a decrease in retrogradely moving APs in all injured axons (Fig. 3F). In contrast, AL dynamics varied depending on axonal outcome: axons which will regenerate or remain static showed reduced retrograde movement, whereas axons which will retract did not differ from pre-injury conditions (Fig. 3G). β1^+^ vesicles became significantly more stationary following axotomy across all conditions (Fig. 3H). Statistical details are provided in Table S2. Due to the high proportion of stationary vesicles after injury, dynamic analysis of LC3⁺-β1⁺ colocalized vesicles was limited and did not yield informative data.

In summary, we characterized vesicle behavior shortly after axotomy and categorized axons based on their regenerative state at 2 h, allowing us to assess whether early vesicle dynamics predict regeneration states. Axotomy induces a rapid and consistent response across axons, characterized by increased numbers of autophagic and integrin vesicles, impaired AP maturation, and an overall decrease in vesicle motility, particularly in the retrograde direction. Only two features differed according to outcome: AL dynamics and the proportion of integrins recycled via autophagy. Axons that later retracted maintained pre-injury AL trafficking, whereas axons that regenerated or remained static showed reduced retrograde transport. In addition, a higher fraction of integrins was recycled by autophagic vesicles in axons that fail to regenerate compared to those that do.

### 4. Autophagic and integrin vesicles responses during regeneration or regeneration failure

At 2 h post-injury, when regenerative outcomes were evident, axonal dynamics were re-evaluated (Fig. 4A; Video S8).

**Figure 4.**
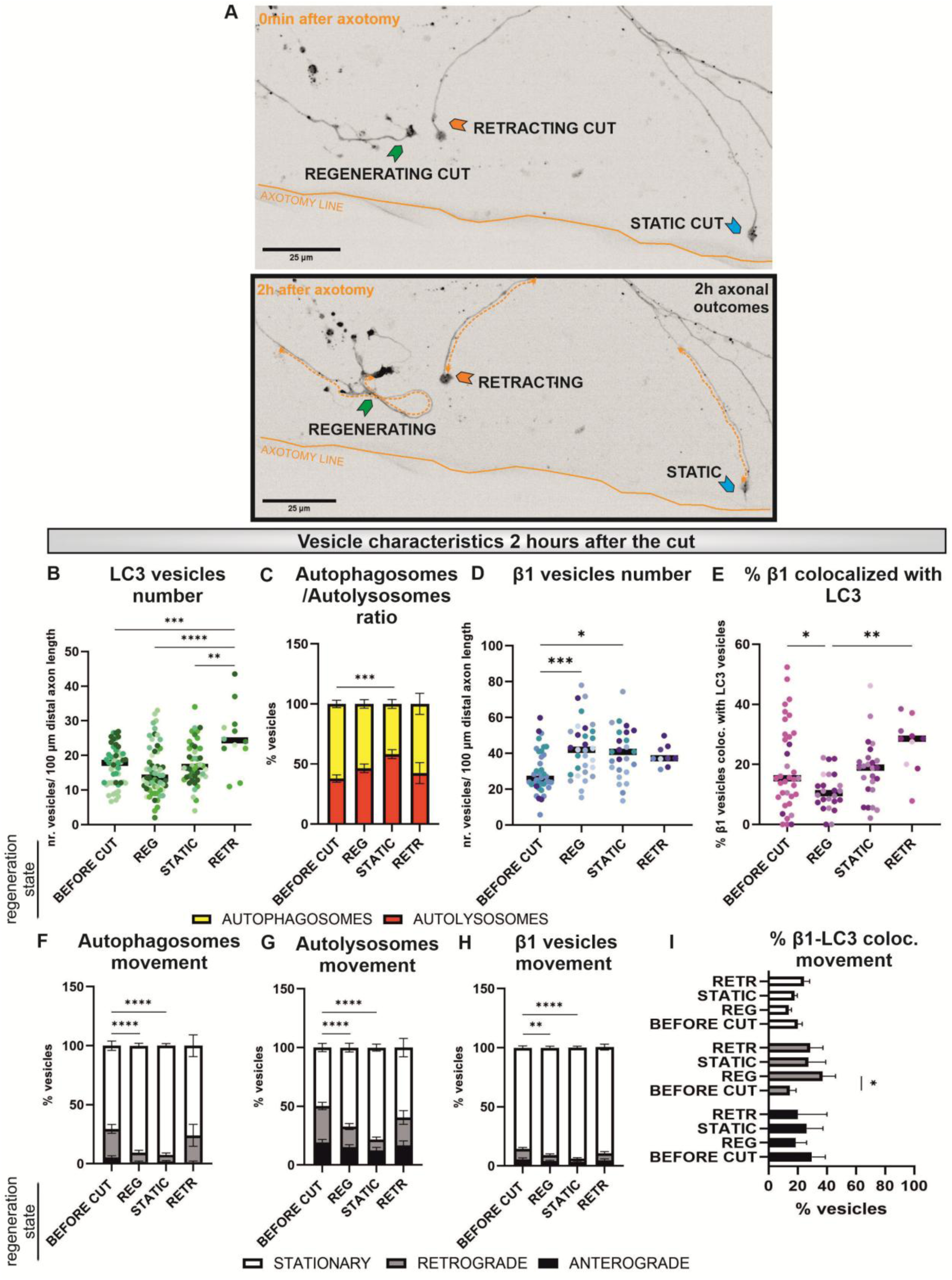
Autophagic and integrins dynamics during regeneration or regeneration failure. (**A**) Representative single-channel (RFP) images from live imaging acquired 2 h after axotomy, focusing on the regeneration phase, with axons classified as regenerated, retracted, or static. Green arrows indicate regenerating axons; blue arrows indicate static axons; orange arrows indicate retracting axons. Orange double arrows indicate the axonal distal segment analyzed. (**B–I**) Quantifications as in Fig. 2. Each dot represents a single axon. Colors indicate independent biological replicates (different animals and imaging days). 8 independent experiments were performed (red+green configuration, n = 60 regenerating axons, n = 46 static axons, n = 12 retracting axons; red+far-red configuration, n = 31 regenerating axons, n = 25 static axons, n = 10 retracting axons). Bars represent mean ± SEM. Statistical analysis was performed using one-way ANOVA (B, D, E) or two-way ANOVA (C, F–I). *p < 0.05, **p < 0.01, ***p < 0.001, ****p < 0.0001.

Regenerating and static axons reduced their autophagic vesicle numbers from the elevated levels observed immediately after injury to values comparable to un-lesioned axons, whereas retracting axons continued to display an increased number (Fig. 4B). Regarding autophagic vesicle composition, only static axons maintained a higher proportion of APs relative to ALs, while both regenerating and retracting axons appeared to restore pre-injury proportions, suggesting a resolution of the injury-induced maturation imbalance (Fig. 4C). The total number of β1^+^ vesicles remained elevated in all injured axons compared to uninjured conditions, with the highest levels observed in regenerating axons (Fig. 4D). LC3⁺-β1⁺ colocalized vesicles followed the same pattern observed immediately after injury; reduced autophagy–integrin colocalization was observed in regenerating axons, whereas increased colocalization was associated with retracting axons (Fig. 4E).

Interestingly, APs, ALs and β1-integrin vesicles remained largely stationary in regenerating and static axons even two hours post-injury, whereas retracting axons have vesicle motility patterns similar to uninjured conditions (Fig. 4F–H). For LC3⁺-β1⁺ colocalized vesicles, we observed a significant increase in retrograde transport in regenerating axons compared to uninjured conditions. This suggests that, although overall vesicle motility is reduced, colocalized autophagy–integrin cargo may undergo a directional shift toward retrograde trafficking during regeneration. We further examined whether the reduction in vesicle motility also occurred within growth cones (Fig. 5A,C; Video S9–S12). The maximum track displacement of LC3⁺ vesicles was reduced in growth cones of regenerating axons compared to uninjured conditions (Fig. 5B) and a similar reduction was observed for β1⁺ vesicles (Fig. 5D). Statistical details are provided in Table S3-4.

**Figure 5.**
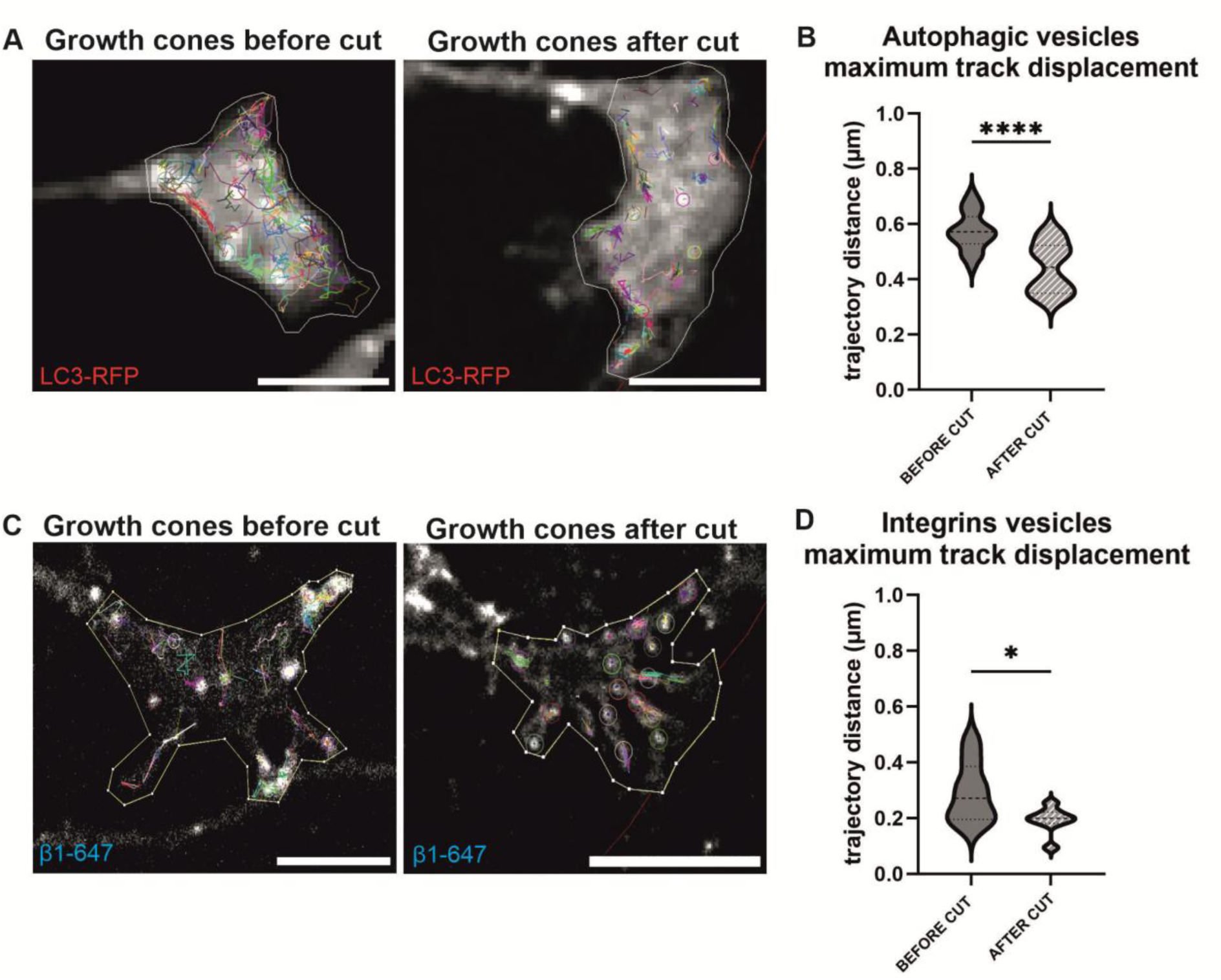
Vesicle motility and trajectory in growth cones before and 2 hours after axotomy. (**A**) Representative trajectories of LC3⁺ vesicles (LC3-RFP) in growth cones before and after injury. Each vesicle track is shown in a different color based on automated TrackMate analysis; zoom-in shows higher-resolution details. (**B**) Quantification of maximum displacement of LC3^+^ vesicles. (**C**) Representative trajectories of β1^+^ vesicles (β1-647). (**D**) Quantification of maximum displacement of β1^+^ vesicles. Statistical analysis was performed using unpaired t-tests. *p < 0.05, ****p < 0.0001. Scale bar, 10 µm.

Together, distinct regeneration states are associated with specific vesicle behaviors. Regenerating axons return to pre-injury levels of total autophagic vesicles while retracting axons retain elevated vesicle numbers. For both regenerating and retracting axons AP:AL ratio returns to pre-injury levels, only axons that remain static continue displaying an increased proportion of APs. The percentage of integrins recycled via autophagy increases in retracting axons, while in regenerating axons decreases, with trafficking biased toward retrograde movement. Interestingly, regenerating axons exhibit predominantly stationary vesicles retained within the distal axon, whereas retracting axons are the only condition in which vesicle trafficking more closely resembles pre-injury dynamics rather than displaying a stationary bias.

Importantly, these data highlight distinct trafficking behaviors between pre-injury growing axons and post-injury regenerating axons and their growth cones.

### 5. Integrins regulate autophagy

Having established that autophagy traffics and recycles a proportion of integrins, we next asked whether a bidirectional relationship exists between these pathways, and integrins in turn, might regulate autophagy. To address this, we pharmacologically manipulated integrin activation and signaling using aggrecan as an integrin inactivator and manganese (Mn²⁺) as an integrin activator [27] and assessed autophagy dynamics (Fig. 6A; Video S13–S16).

**Figure 6.**
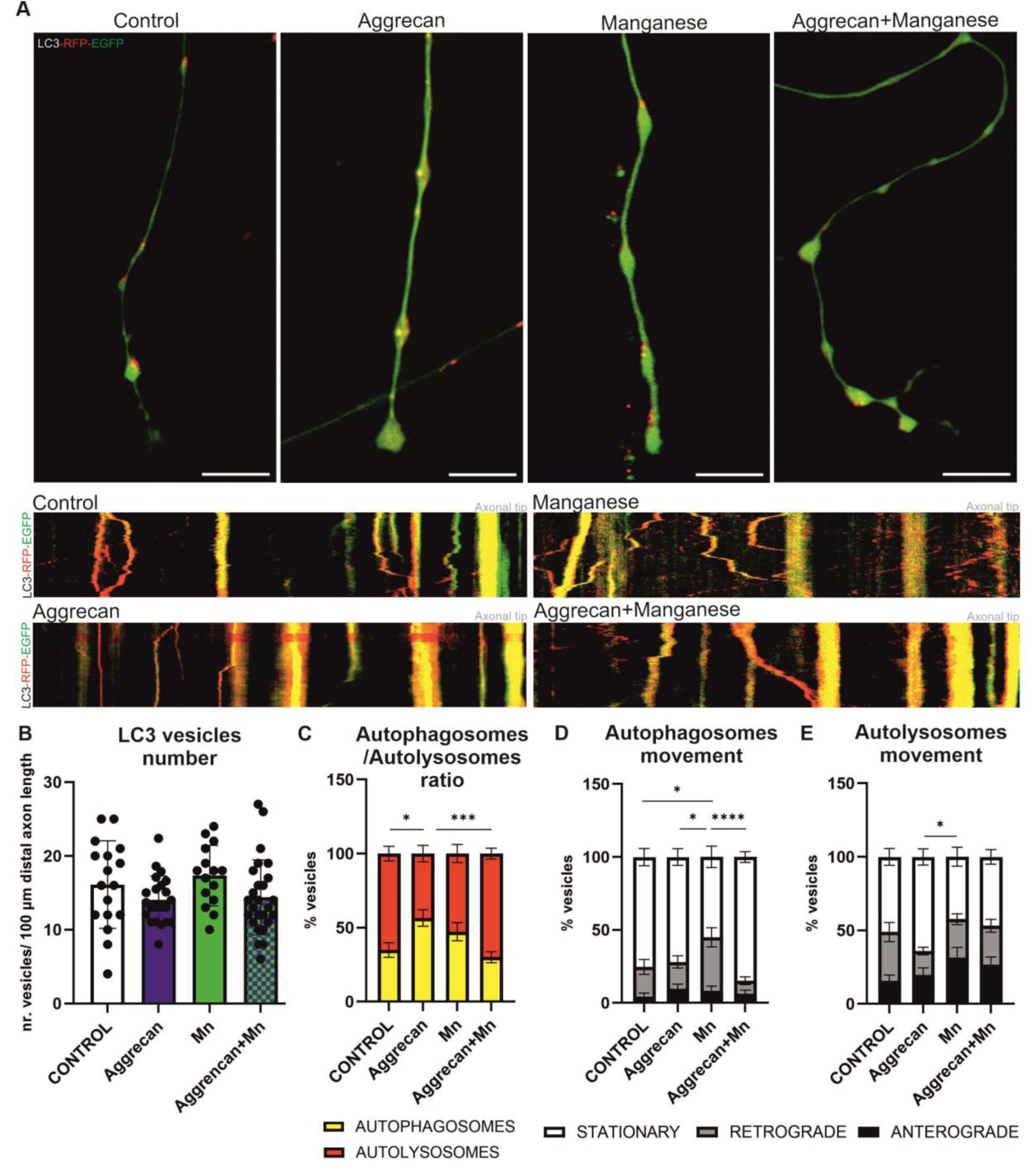
LC3-RFP-EGFP live imaging in typical axons following integrin manipulation. (**A**) Representative distal axons and corresponding kymographs under control, aggrecan, Mn²⁺, and combined aggrecan + Mn²⁺ conditions. (**B**) Total number of LC3^+^ vesicles. (**C**) Proportion of AP:AL. (**D– E**) Directionality analysis for APs (**D**) and ALs (**E**). Each dot represents a single axon. 3 independent experiments were performed (n = 17 control, n = 22 aggrecan, n = 15 Mn²⁺, n=26 aggrecan + Mn²⁺). Bars represent mean ± SEM. Statistical analysis was performed using one-way ANOVA (B) or two-way ANOVA (C–E). *p < 0.05, ***p < 0.001, ****p < 0.0001. Scale bar, 10 µm.

The results indicate that integrin modulation does not affect the total number of autophagic vesicles (Fig. 6B). However, integrins appear to contribute to AP maturation: inhibition of integrins with aggrecan shifted the balance toward increased APs relative to ALs, whereas co-application of Mn²⁺ restored the proportion to control levels (Fig. 6C). Integrin signaling also influenced autophagic vesicle trafficking: activation with Mn²⁺ reduced the fraction of stationary APs, an effect that was abolished by co-treatment with aggrecan (Fig. 6D), and integrin inhibition with aggrecan slightly increased the proportion of stationary ALs, although this effect reached significance only when compared to Mn²⁺-treated conditions. Notably, Mn²⁺ treatment also increased the proportion of ALs undergoing anterograde transport (control: 15%; Mn²⁺: 31%), although this did not reach statistical significance (Fig. 6E). Statistical details are provided in Table S5.

In summary, while autophagy contributes to integrin recycling (as shown above), this data suggests a reciprocal regulatory role of integrins in autophagy. Specifically, integrin inhibition with aggrecan increases the AP:AL ratio, whereas integrin activation with Mn²⁺ enhances retrograde transport of APs.

### 6. Effect of 3-methyladenine, berbamine and rapamycin treatment on axonal autophagy dynamics

We treated DRG explants with three pharmaceuticals that have been widely used for acute manipulation of autophagy: 3-methyladenine (3-MA) to decrease the biogenesis of APs [28], berbamine (BBM) to block the maturation of APs to ALs [29], and rapamycin (RAPA) to increase autophagy through its action on mTOR [30]. We corroborated the effects of these chemicals on APs and ALs using both immunocytochemistry (Fig. 7A) and live-imaging analysis (Fig. 8A). The immunocytochemistry results demonstrate that the chemicals produced the expected effects in a dose-dependent manner. 3-MA decreased the total number of vesicles (Fig. 7B) and did not significantly affect the proportion of AP:AL, indicating no effect on AP maturation (Fig. 7E). BBM treatment at 10 µM caused an increase in the total number of vesicles as observed by immunocytochemistry (Fig. 7C), probably due to AP accumulation, as they fail to convert to AL, leading to a significant increase in APs (Fig. 7F). RAPA increased the total number of vesicles at concentrations as low as 1 nM (Fig. 7D), suggesting an upregulation of AP generation with normal maturation. At 1 nM, RAPA did not affect the proportion of AP:AL but at 100 nM concentration there was a significant increase in ALs, suggesting enhanced AP-lysosomes fusion (Fig 7G). Live imaging analysis was consistent with these findings. Treatment with 2.5 mM of 3-MA confirmed a decrease in total number of vesicles with no significant changes in vesicle proportions compared to the control (Video S17-18). BBM treatment did not show an increase in total number of autophagic vesicles but caused AP maturation disruption, leading to altered vesicle proportions (Video S19). RAPA treatment live-imaging analysis resulted in a clear increase in total number of vesicles and a tendency towards an increase in ALs (Video S20). Statistical details are provided in Table S6-7. In summary, 3-MA effectively decreased the total number of vesicles, BBM blocked AP maturation, and RAPA increased the number of autophagic vesicles, corroborating that the chemicals have the desired effect (Fig. 8B,C).

**Figure 7.**
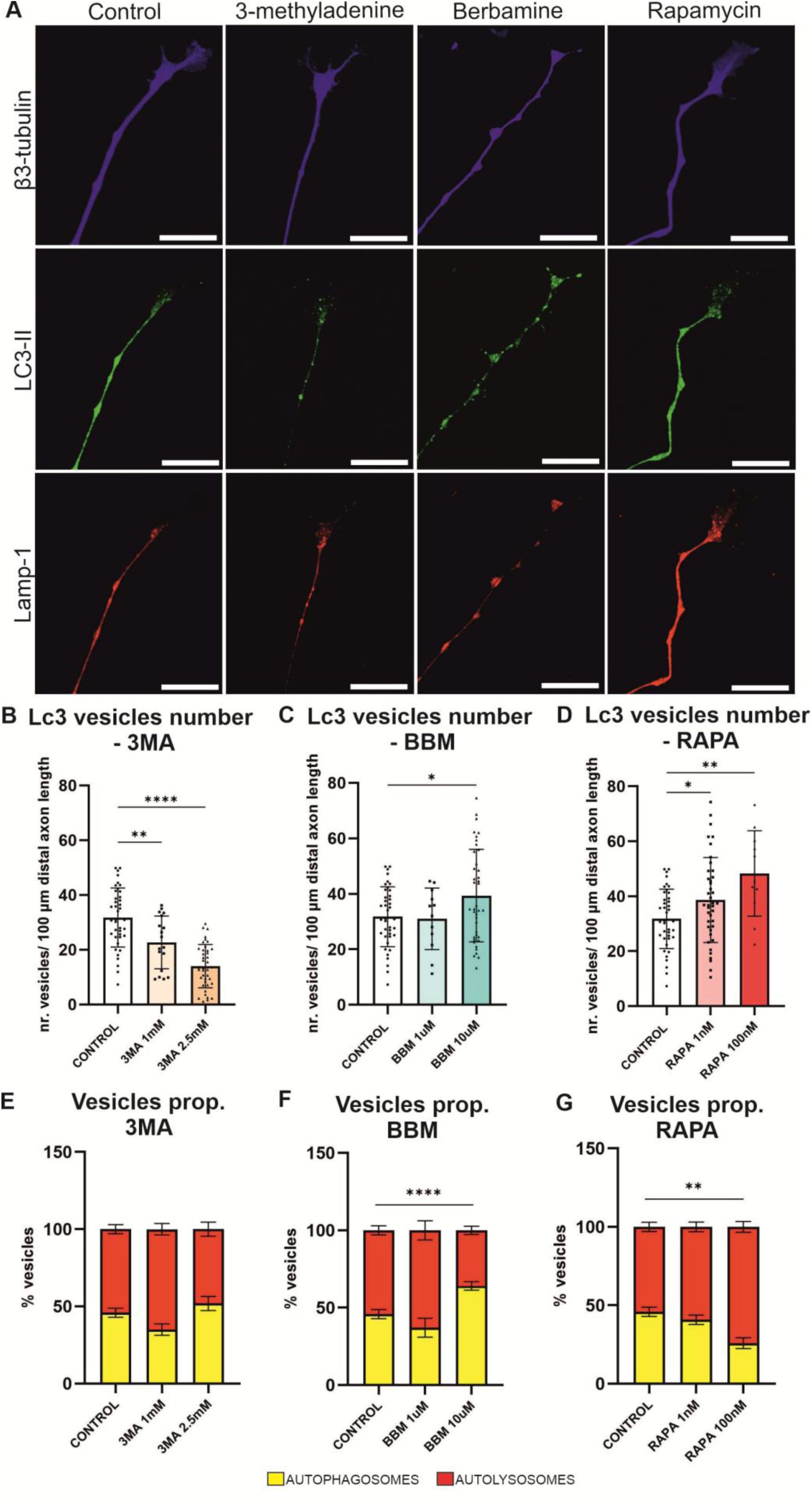
Immunocytochemical assessment of autophagic vesicles following chemical manipulation. (**A**) Representative images of distal axons treated with 3-MA, BBM, or RAPA). (B–D) Quantification of LC3-II^+^ vesicles across increasing concentrations of 3-MA (**B**), BBM (**C**), and RAPA (**D**). (**E–G**) Proportion of APs (LC3-II^+^ LAMP1^-^) and ALs (LC3-II^+^-LAMP1^+^) across increasing concentrations of 3-MA (**E**), BBM (**F**), and RAPA (**G**). Each dot represents a single axon. 5 independent experiments were performed. Bars represent mean ± SD (B–D) or mean ± SEM (E–G). Statistical analysis was performed using one-way ANOVA (B–D) or two-way ANOVA (E–G). *p < 0.05, **p < 0.01, ****p < 0.0001. Scale bar, 25 µm.

**Figure 8.**
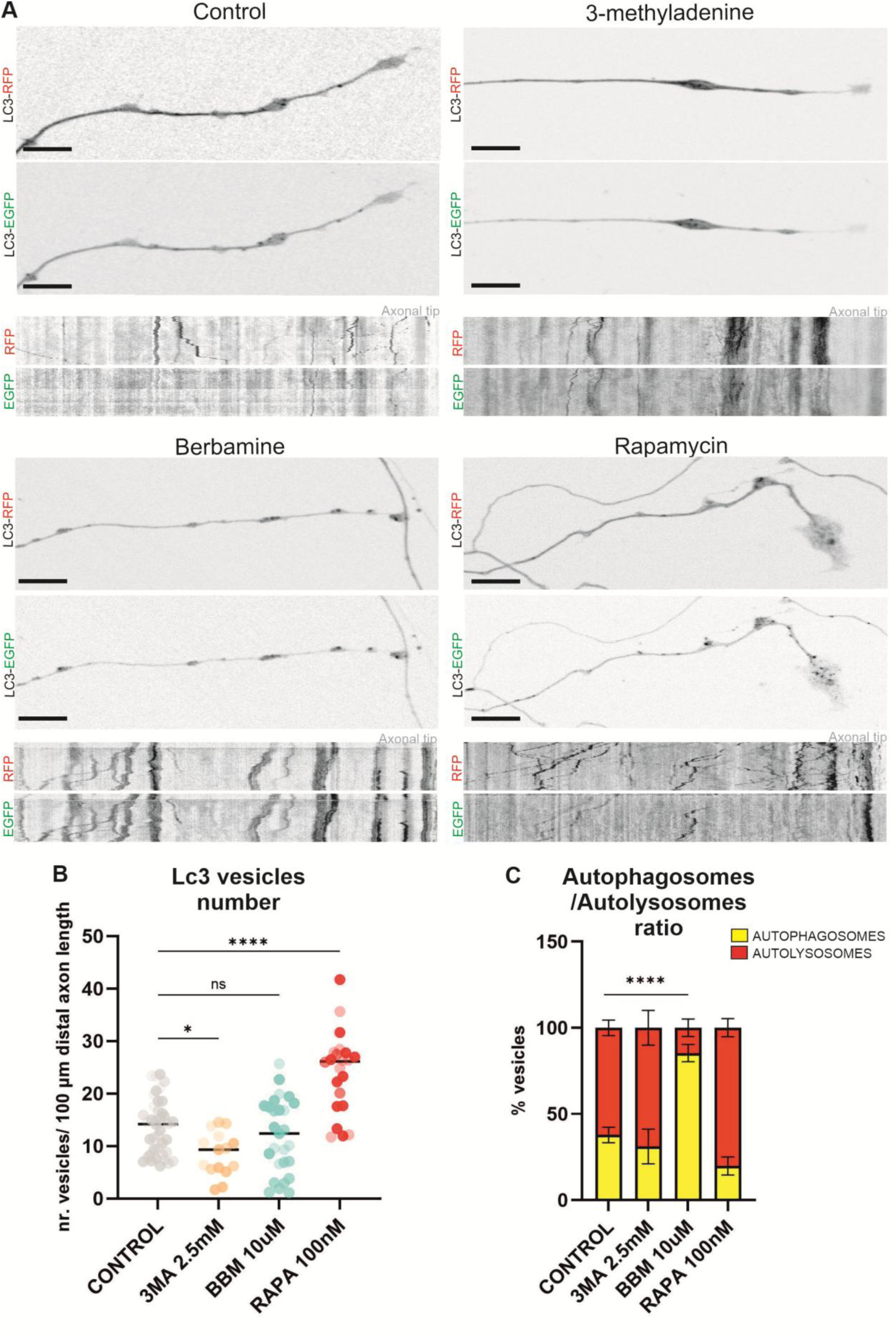
Live imaging assessment of autophagic vesicles following chemical manipulation. (**A**) Representative live imaging frames of distal axons treated with 3-MA, BBM, or RAPA. (**B**) Total number of LC3^+^ vesicles. (**C**) Proportion of AP:AL. Each dot represents an individual axon (n = 40 control, n = 15 3-MA, n = 27 BBM, n = 22 RAPA); colors shade denotes independent experiments (n = 3). Bars represent mean ± SEM. Statistical analysis was performed using one-way ANOVA (B) or two-way ANOVA (C). *p < 0.05, ****p < 0.0001. Scale bar, 10 µm.

### 7. Autophagy manipulation has effects on normal axon growth and regeneration

We treated adult DRG explants surrounded by a halo of axons with 2.5 mM 3-MA, 10 µM BBM, or 100 nM RAPA. Three hours following the start of the treatment, explants were either left intact (Fig. 9A) or subjected to axotomy (Fig. 9B). Axons were imaged with phase-contrast microscope, returned to the incubator for two hours, and then re-imaged to assess growth and regeneration.

**Figure 9.**
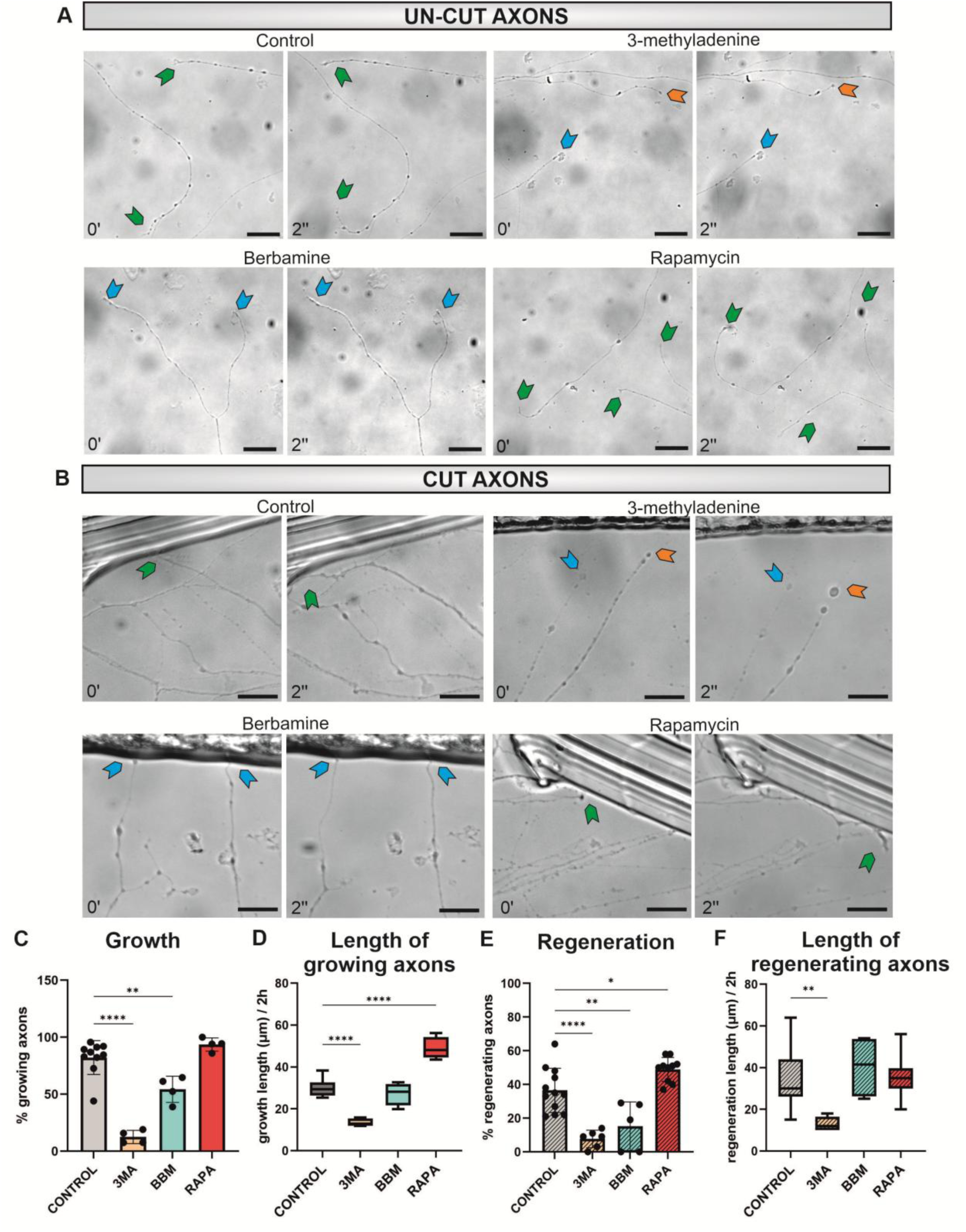
Axonal growth and regeneration following autophagy modulation. Treatment was applied 3 h before analysis. (**A**) Representative bright-field images of axonal growth over 2 h. (**B**) Representative images of axonal regeneration after axotomy over 2 h. Green arrows indicate growing/regenerating axons; blue arrows indicate static axons; orange arrows indicate retracting axons. (**C**) Percentage of growing axons. (**D**) Axon growth length increase during 2 h. (**E**) Percentage of regenerating axons. (**F**) Regenerated axon length during 2 h. Each dot represents an independent experiment (n = 12 control, n = 6 3-MA, n = 5 BBM, n = 10 RAPA). Bars represent mean ± SD. Statistical analysis was performed using one-way ANOVA. *p0.05, **p < 0.01, ****p < 0.0001. Scale bar, 25 µm.

Under physiological conditions, approximately 80% of axons exhibited growth over the two-hour period, with an average elongation of ∼30 µm. Following axotomy, 40% of axons initiated regeneration, with an average regenerative outgrowth of ∼35 µm over the same time window. Pharmacological inhibition of autophagy with 3-MA strongly impaired both processes, reducing the proportion of growing and regenerating axons and significantly decreasing axonal elongation in both contexts. In contrast, inhibition of AP maturation with BBM reduced the proportion of axons that grew and regenerated but did not affect their elongation rate. Conversely, enhancing autophagy with RAPA increased the proportion of axons undergoing growth and regeneration. However, RAPA significantly increased axonal elongation under physiological conditions, reaching an average of 50 µm over two hours but did not affect regenerative outgrowth length (35 µm) (Fig. 9C–F). Statistical details are provided in Table S8.

In summary, the results support a major role for autophagy in axon growth and regeneration, and its modulation significantly influences regenerative outcomes. Reducing autophagy or blocking AP maturation impaired both processes, whereas increasing autophagy enhanced basal growth and increased the proportion of regenerating axons.

### 8. Rapamycin effects on integrin recycling

RAPA enhanced both axonal growth and regeneration. To further investigate the underlying mechanisms, we performed live imaging of β1 integrin trafficking in axons following 3 hours of RAPA treatment (Fig. 10A, Video S21). As expected, RAPA increased the total number of LC3⁺ vesicles (Fig. 10B). We also observed a significant increase in β1⁺ vesicles (Fig. 10C) and a decrease in the percentage of β1⁺ vesicles that colocalize with LC3⁺ vesicles, indicating lower recycling of integrins via autophagy (Fig. 10D). A significantly higher fraction of β1⁺ vesicles remained stationary under RAPA treatment (Fig. 10E). In addition, the number of colocalized LC3⁺-β1⁺ vesicles undergoing retrograde transport was markedly increased (10F). Statistical details are provided in Table S9. Notably, this combination of increased integrin vesicle abundance, increased integrin-vesicle stalling at the distal axon, reduced autophagy– integrin colocalization, and enhanced retrograde transport of colocalized vesicles resembles the vesicle dynamics observed in regenerating axons (Fig. 4).

**Figure 10.**
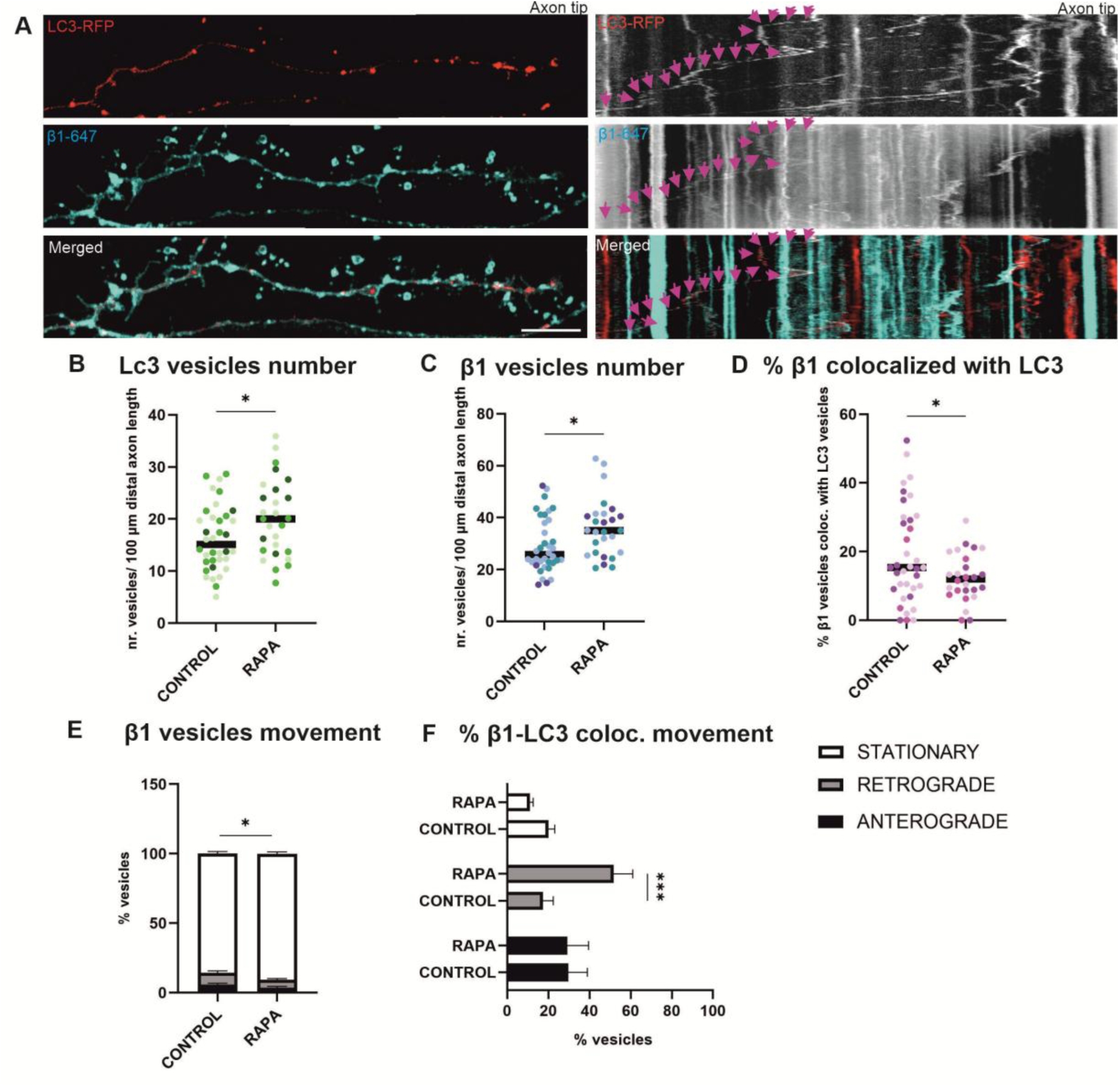
Further analysis of rapamycin effects on integrin vesicle dynamics. (**A**) Representative livimaging frames and corresponding kymographs of RAPA-treated axons showing LC3 (RFP) and β1 integrin (far-red) vesicles. Pink arrows denote motile LC3-β1-integrin colocalized vesicle. (**B–D**) Quantification of LC3^+^ vesicles (**B**), β1^+^ vesicles (**C**), and their colocalization (**D**). (**E–F**) Directionality analysis for β1 vesicles (**E**), and β1^+^-LC3^+^ vesicles (**F**). Each dot represents an individual axon (n = 36 control, n = 27 RAPA); colors denote independent experiments (n = 3). Bars represent mean ± SEM. Statistical analysis was performed using unpaired t-tests (B-D) or two-way ANOVA (E,F). *p < 0.05, **p < 0.01, ***p < 0.001. Scale bar, 10 µm.

In summary, RAPA produced the expected effect on autophagic vesicle behavior and additionally altered β1 integrin distribution and trafficking at the distal axon. Together, these findings suggest that RAPA primes axons toward a regeneration-like state, which is unlikely to arise solely from increased autophagic vesicle biogenesis. Instead, RAPA appears to modulate autophagy–integrin coupling and vesicle trafficking dynamics, including cargo association and transport directionality.

## DISCUSSION

Axon regeneration requires coordinated and tightly regulated changes in neuronal homeostasis, including membrane resealing, regulation of Ca²⁺ and reactive oxygen species (ROS) influx, cytoskeletal remodeling, and the localized translation and transport of signaling and structural molecules to and from the injury site, alongside retrograde injury signaling and sustained metabolic demand [20]. Following axonal injury, neurons first form a retraction bulb that stabilizes the damaged membrane, after which successful regeneration proceeds through the emergence of a new growth cone and axonal elongation.

Integrins are key regulators of axonal growth, mediating dynamic adhesion to the extracellular matrix (ECM) through coordinated cycles of attachment and detachment that drive neurite extension. Integrin-based approaches have emerged among the most successful experimental therapeutic targets for adult axon regeneration. Restoring expression of the tenascin receptor α9 integrin, particularly when combined with the integrin activator kindlin-1, promotes robust regeneration of adult sensory axons through spinal cord lesions and supports long-distance growth extending to the medulla, accompanied by functional recovery [15–17]. In addition to sensory neurons, manipulation of integrin signaling, activation, or trafficking enhances neurite growth and regenerative responses in adult retinal ganglion cells [31] and cortical neurons [32], highlighting integrins as broadly relevant regulators of neuronal repair [11,33].

Emerging evidence from non-neuronal systems and from developmental and neurodegenerative contexts suggests a functional interplay between autophagy and adhesion turnover [19,34]. Whether such coupling also contributes to axonal regeneration remains largely unexplored. More broadly, the role of autophagy in regeneration is still debated [35]. Here, we propose that a key determinant of autophagy function in axon regeneration is not its overall level, but rather its cargo selectivity and the highly compartmentalized nature of neuronal trafficking.

### 1. Autophagic vesicles and integrins in growth cones

Although autophagy has gained increasing attention in axons, its characterization within growth cones remains incompletely described. We find that adult DRG growth cones contain a high abundance of autophagic vesicles, with APs clustered in the central domain while ALs can be find all the way to the leading edge of membrane remodeling in filopodia (Fig. 1A). Although lysosomal transport to growth cones has been implicated in axon extension and growth cone dynamics [36,37], the spatial distribution of ALs within adult growth cones has received little attention. Our observations suggest that autophagic degradation is strategically positioned near sites of growth cone extension, potentially supporting local recycling processes required for growth cone motility (actin turnover, local energy balance, etc.). Whether these autophagic structures arise locally within growth cones or are delivered from more distal axonal compartments remains an open question. Notably, we frequently observe LC3⁺ vesicles in close spatial proximity to β1⁺ vesicles, displaying coordinated movement (Fig. 1C). β1-integrin is predominantly trafficked through Rab11-positive recycling endosomes [14], a compartment that has been implicated in supplying membrane and regulatory components for AP biogenesis in specific cellular contexts [38]. These observations suggest an interaction between recycling endosomal trafficking and autophagic pathways within growth cones, consistent with previously reported crosstalk between these systems.

However, we observed only small amounts of actual colocalization between LC3⁺ and β1⁺ vesicles, restricted to the growth cone central domain (Fig. 1C). Instead, transient colocalization occurred predominantly within wave-like membrane remodeling events at the leading edge of growth cones. Here, surface patches of β1-integrin signal preceded LC3 recruitment, followed by brief colocalization and dissipation of both signals (Fig. 1D). This behavior is consistent with a role of LC3 in FA turnover, as described in FA-phagy, where LC3 associates with adhesion components such as paxillin through selective receptors as NBR1, c-Cbl, and SRC [18]. Neuronal growth cones form transient focal adhesions with extremely fast assembly-disassembly cycles which explain the “pulses” of wave-like remodeling events we observed. However, the limited overlap between vesicular LC3 and β1-integrin suggests that autophagy is unlikely to represent the major pathway for bulk integrin turnover within growth cones. Instead, and in agreement with previous studies, integrin trafficking in this compartment is likely mediated predominantly through endosomal Rab GTPase-dependent recycling pathways [14].

Regarding vesicle dynamics, autophagic vesicle movement within growth cones differs markedly from that observed in distal axons. In growth cones, autophagic vesicles exhibit a lack of directionality, frequent pausing, and confined, exploratory movements, whereas in distal axons trafficking is more directional and sustained. One of the main findings of this study is that mechanical injury induces both acute and longer-term alterations in intracellular trafficking dynamics. Accordingly, axons under physiological growth conditions display distinct trafficking behavior compared to regenerating axons following injury (discussed in the following section), a distinction that is also reflected at the level of their respective growth cones. At 2 h post-injury, successfully regenerated growth cones exhibited a reduction in trafficking dynamics, reflected by decreased total track displacement of both LC3⁺ and β1⁺ vesicles, compared with growth cones of uninjured axons (Fig. 5). Together, these data suggest that although axonal growth and regeneration share core mechanisms, they also engage distinct regulatory programs. In particular, vesicle trafficking dynamics in distal axons and growth cones appear to be differentially modulated depending on the physiological versus regenerative state. This reduction in motility may reflect a shift toward increased spatial confinement and stabilization of vesicular pools during early regenerative stages.

### 2. Autophagy–integrin coupling during axonal remodeling

Using live imaging across injury, growth, and regeneration paradigms (Figs. 1–5), we identify distinct intracellular signatures which correspond with different axonal outcomes.

#### **a)** Autophagic vesicle abundance and maturation state

Under basal conditions, axons can either grow or remain static. While these two populations did not differ in total autophagic vesicle number (Fig. 2B), static axons exhibited a significantly increased ratio of AP:AL, suggesting impaired AP maturation (Fig. 2C). Exposure to inhibitory extracellular cues such as chondroitin sulfate proteoglycans (CSPGs) induced a comparable increase in the AP:AL ratio, further linking impaired AP maturation to growth failure [8,39]. Similarly, disruption of integrin function by aggrecan application, a condition known to inhibit axon growth, was associated with a comparable increase in the AP:AL ratio (Fig. 6C). Pharmacological inhibition of AP–lysosome fusion using BBM (Fig. 7F, 8C) resulted in pronounced axonal stalling (Fig. 9C). Together, all these findings suggest that efficient AP maturation is associated with a growth-permissive axonal state, whereas impaired maturation is consistently linked to axonal growth arrest across multiple experimental conditions.

At an early stage following axotomy, all severed axons exhibit similar morphological features, including retraction bulb formation and growth arrest, independent of their eventual outcome (Fig. 3A). At this point, axons are rapidly engaging a coordinated intracellular response involving damage-associated molecular patterns (DAMPs), ROS, and increased intracellular Ca²⁺, all of which are known to activate autophagy. Previous studies have reported transcriptional upregulation of autophagy-related genes following neuronal injury, suggesting activation of autophagy as a stress-responsive pathway [40]. However, accumulating evidence also indicates that increased autophagy does not necessarily reflect productive autophagic flux, as impaired AP maturation and disrupted degradation have been reported in several injury paradigms [41–43]. In agreement with this, our live imaging data immediately after axotomy revealed increased autophagic vesicle abundance (Fig. 3B) and an elevated AP:AL ratio (Fig. 3C), consistent with impaired autophagic maturation rather than solely increased pathway induction.

At later time points (∼2 h post-injury), distinct vesicles patterns emerged across axonal outcomes (Fig. 4A). In axons that successfully regenerated, both AP:AL ratios and total autophagic vesicle abundance returned to basal pre-injury levels, indicating recovery from the transient injury-induced disruption of autophagic flux. In contrast, retracting axons failed to normalize autophagic vesicle abundance, which remained elevated, whereas static axons retained a persistent AP:AL imbalance indicative of impaired maturation (Fig. 4 B,C). These data suggest that axotomy induces a transient but uniform disruption of autophagic homeostasis across all severed axons, characterized by increased autophagic vesicles and defective maturation. Successful regeneration is associated with a return to basal autophagy levels and maturation state, whereas failure to regenerate correlates with persistent deviation from this baseline, either through sustained vesicles increase or continued maturation defects.

#### **b)** Integrins and their autophagy-mediated turnover

LC3-β1 colocalization showed a striking spatial segregation, being minimal in growth cones but substantial in distal axons.

Under basal conditions, growing and static axons showed similar numbers of β1-integrin vesicles and comparable levels of LC3-β1 colocalization (Fig. 2D, E), with differences instead emerging at the level of vesicle trafficking dynamics, which is discussed in the following section.

Following axotomy, all injured axons exhibited increased numbers of β1-integrin-containing vesicles (Fig. 3D). This observation is consistent with previous studies demonstrating that axonal injury enhances integrin availability at the injured axon through coordinated changes in transcription, trafficking and subcellular redistribution mechanisms that support injury response and regeneration [11,16]. Notably, although both autophagic vesicles and β1-integrin vesicles increased in all injured axons, LC3-β1 colocalization was differentially regulated depending on regenerative outcome. Axons that remained static or retracted showed increased LC3-β1 association, whereas regenerating axons displayed reduced colocalization compared with uninjured controls (Fig. 3,4 E). These findings suggest that autophagy-associated integrin turnover is differentially regulated during successful regeneration and failed regenerative responses. Importantly, this behavior differs from that observed during physiological axon growth, highlighting again that regeneration is not simply a recapitulation of developmental or basal growth programs.

Studies in epithelial cells, fibroblasts and cancer cells have shown that autophagy contributes to FA remodeling through the turnover of integrins and adhesion-associated proteins such as paxillin. However, the effects of autophagy on motility are context dependent, as both insufficient and excessive autophagic processing of adhesion complexes can impair migration [24,44–46]. Growth cones similarly require a finely balanced level of integrin-mediated adhesion to generate traction while maintaining sufficient adhesion turnover for forward advance. Increased LC3-β1 association in retracting axons may therefore reflect enhanced autophagic processing of integrin-containing compartments, potentially limiting the adhesive interactions required for regeneration. Conversely, reduced LC3-β1 association in regenerating axons may help maintain integrin availability and support productive growth. Nevertheless, whether these interactions represent degradation, recycling, storage, or redistribution of integrins remains to be determined.

#### **c)** Vesicle trafficking and injury-induced transport reorganization

Vesicle trafficking represented the most pronounced alteration in our dataset. Under physiological conditions, non-growing static axons exhibited reduced retrograde movement of APs, whereas AL motility was largely preserved (Fig. 2F, G), suggesting that these vesicle populations are differentially regulated, consistent with previous reports [47]. In the same axons, β1-integrin–containing vesicles showed a predominance of stationary vesicles (Fig. 2H). Notably, although both APs and β1-integrin vesicles were largely immobile, LC3-β1 colocalized vesicles showed enrichment in the retrograde fraction and were absent from anterograde transport (Fig. 2I). The divergence between independently tracked vesicle populations, consistent with previous reports [48], suggests an active reorganization of autophagy-associated integrin handling rather than a global transport failure.

Immediately after injury, vesicle motility was broadly suppressed across APs, ALs, and β1-integrin vesicles, which became predominantly stationary (Fig. 3F, G, H). However, the preservation of AL movement in axons which will eventually retract, argues against a purely mechanical disruption of microtubule-based transport. Instead, these data support a model in which vesicle stalling at distal axons represents a regulated response to injury.

At later stages, regenerating axons maintained a higher proportion of stationary vesicles compared with uninjured axons, whereas retracting axons reverted to pre-injury trafficking patterns (Fig. 4F, G, H). This suggests that injury induces a switch in trafficking programs that promotes local retention of vesicular cargo, potentially supporting membrane repair and growth cone formation. A notable feature of regenerating axons was the increased proportion of LC3-associated β1-integrin vesicles within the retrograde population, despite their overall low abundance and predominantly stationary state (Fig. 4I). This observation is noteworthy because retrograde trafficking has been linked to axon extension, although the specific mechanisms and cargoes responsible remain incompletely understood [49].

Collectively, these findings support a model in which axonal injury triggers a coordinated reorganization of vesicle dynamics characterized by reduced global mobility and selective stabilization of vesicle pools. This reorganization differs across regenerative outcomes and is consistent with previous studies showing that injury alters axonal transport, organelle distribution, and cytoskeletal organization [50–52]. Here, we extend these observations by demonstrating outcome-specific trafficking signatures in the distal axon, suggesting that vesicle retention may be a key feature of successful regeneration.

In conclusion, our live imaging across multiple time points and experimental conditions reveals distinct intracellular signatures associated with different axonal states. Uninjured axons display trafficking dynamics that are clearly distinct from those of injured axons, while growing, static, regenerating, and retracting axons each exhibit specific vesicular transport patterns. Across these conditions, the most consistent differences are observed in vesicle trafficking dynamics rather than vesicle abundance alone, highlighting the importance of assessing both spatial and temporal organization of intracellular transport. These findings underscore the value of live imaging approaches to capture dynamic, compartment-specific behavior, particularly in highly polarized and structurally compartmentalized cells such as neurons.

### 3. Integrin signaling regulates autophagy

The relationship between integrin signaling and autophagy remains relatively underexplored, particularly in the nervous system, with most studies conducted in non-neuronal cell types. Current evidence suggests a bidirectional interplay between these pathways, whereby loss of integrin-mediated adhesion induces autophagy, while integrin activation generally suppresses autophagic activity [25,53,54]. In contrast, our findings indicate that integrin signaling does not regulate AP biogenesis in neurons (Fig. 6B), but rather influences later stages of the pathway, specifically AP maturation and trafficking.

Aggrecan is a major CSPG of the ECM and perineuronal nets that contributes to the inhibitory environment of the glial scar, thereby limiting axon regeneration after injury [55]. Aggrecan inhibits integrin signaling and as a result, growth cones lose traction on the substrate and axons stall in CSPG-rich environments [27]. Interestingly, Sakamoto et al. previously demonstrated that CSPGs impair axonal autophagy and growth through activation of the receptor-type protein tyrosine phosphatase PTPσ, leading to defects in AP maturation and trafficking [8]. However, integrin signaling was not investigated in that study. Our findings with aggrecan also showed impaired maturation and trafficking (Fig. 6C, E), suggesting that CSPGs inhibit axonal growth and regeneration through converging mechanisms involving both PTPσ-dependent signaling and impaired integrin–ECM interactions.

Conversely, activation of integrins with manganese Mn²⁺, a divalent cation that stabilizes integrins in a high-affinity conformation and promotes axon growth in DRG neurons [11,27], enhanced AP retrograde transport and showed a strong trend towards increased anterograde AL transport (Fig. 6D, E). These findings support a role for integrin signaling in regulating autophagic trafficking dynamics. However, Mn²⁺ exposure has recently been reported to disrupt AP maturation and lysosomal degradation in hippocampal neurons, leading to synaptic dysfunction [56]. This apparent discrepancy suggests that the effects of integrin activation on intracellular trafficking are highly context-dependent and may vary across neuronal populations and experimental paradigms.

Collectively, our data support the existence of a bidirectional relationship between ECM– integrin signalling and autophagy. While ECM cues regulate AP maturation and trafficking through integrin-dependent mechanisms, autophagy in turn influences the turnover and availability of integrins.

### 4. Rapamycin primes axons for regeneration by altering autophagy dependent integrin recycling

To further investigate the role of autophagy in axon growth and regeneration, explants were treated with pharmacological modulators targeting distinct stages of the autophagic pathway.

The use of pharmacological tools allowed us to study immediate effects of autophagy modulation. 3-MA was used to inhibit AP biogenesis by blocking class III phosphoinositide 3-kinases (PI3Ks), which are required for omegasome formation and early AP initiation at the endoplasmic reticulum [28]. To interfere with AP maturation, BBM was applied, disrupting SNAP29–VAMP8 interactions and thereby inhibiting SNARE-dependent fusion between AP and AL, without directly affecting lysosomal function [29]. Autophagy was enhanced using RAPA, an inhibitor of mTORC1, central negative regulator of autophagy [30].

Dose-dependent effects of these compounds were first validated using immunocytochemistry and live imaging (Figs. 7–8). Subsequent treatment of explants revealed that inhibition of autophagy at either the biogenesis or maturation stage impaired axonal growth and regenerative capacity, whereas autophagy enhancement promoted them (Fig. 9). These results are consistent with previous studies highlighting autophagy as a key process supporting axonal extension and neuroprotection, while defects in AP formation or maturation compromise neuronal survival and regenerative potential [57,58].

We performed live imaging under RAPA treatment while simultaneously monitoring integrin dynamics. RAPA not only increased the number of autophagic vesicles (Fig. 10B) but also elevated the number of integrin-positive vesicles (Fig. 10C) and altered their trafficking and recycling. Specifically, integrin-containing vesicles accumulated in distal axons (Fig. 10E) with reduced autophagy-recycling (Fig. 10D), while retrograde transport of colocalized vesicular structures was enhanced (Fig. 10F), a pattern that closely resembled that observed during regenerative conditions (Fig. 4 D, E, H, I). These findings suggest that RAPA promotes a regeneration-competent state not simply through global upregulation of autophagy, but through coordinated changes in autophagy–integrin coupling and trafficking dynamics. RAPA-induced regulation of integrin expression and trafficking has been reported in other systems, supporting this interpretation [59]. Importantly, these results help reconcile apparently conflicting roles of autophagy in axon regeneration by emphasizing that functional outcomes depend not only on overall autophagic activity, but also on flux directionality, subcellular compartmentalization, and cargo specificity.

In conclusion, our findings identify a previously unrecognized reciprocal relationship between integrin trafficking and autophagy in adult sensory axons, thereby linking ECM cues with intracellular homeostasis. Autophagic vesicle dynamics correlate with axonal growth, regeneration, and retraction, and trafficking, maturation, and cargo selectivity of autophagic vesicles, rather than autophagy induction per se, underlies the pro-regenerative effects of rapamycin. While the molecular mechanisms coordinating these processes remain to be elucidated, and *in vivo* validation is currently limited by the technical challenges of imaging autophagic vesicle dynamics in the intact nervous system, our study provides a new conceptual framework for understanding how autophagy contributes to axon regeneration and highlights cargo-specific autophagic trafficking as a potential therapeutic target.

## MATERIALS AND METHODS

### DRG explants

Adult three months old WT C57BL/6 and C57BL/6-Tg(CAG-RFP/EGFP/Map1lc3b)1Hill/J (Jackson Laboratory) mice were fully anesthetized with isoflurane, decapitated, and dorsal root ganglia (DRGs) were collected. All experiments were performed in accordance with the European Communities council directive of 22nd of September 2010 (2010/63/EU), follow the ARRIVE guidelines (https://arriveguidelines.org/) and were approved by the Ethics Committee of the Institute of Experimental Medicine ASCR, Prague, Czech Republic. Each DRG was deprived of the meningeal membrane and trimmed into two-four pieces. Explants were plated on µ-Slide Angiogenesis chambers (Ibidi, 81506) previously treated with PDL overnight (Sigma-Aldrich, P1149; 20 µg/mL) and laminin (Sigma-Aldrich, L2020; 10 µg/mL) for two hours. Explants were maintained three days *in vitro* (3DIV) with daily media refresh, which consisted of: ITS (Sigma-Aldrich, I2521; 1%), PSF (Sigma-Aldrich, P4333; 2%), NGF (Sigma-Aldrich, N8133; 10 ng/mL) and Mitomycin-C (Sigma-Aldrich, M4287; 1 µg/mL) in DMEM (Thermofisher, 10566016).

### Chemicals manipulation

On day *in vitro* 3 (DIV3), chemical manipulation experiments were performed. To modulate autophagy, the following compounds were used: 3-methyladenine (3-MA, Abcam, ab120841), rapamycin (RAPA, ChemScene, 53123-88-9), and berbamine (BBM, Sigma-Aldrich, 547190). Cells were treated with 3-MA and RAPA for 3 hours prior to analysis, whereas BBM was applied overnight. Each compound was tested at multiple concentrations to evaluate dose-dependent effects, and the optimal concentration (2.5 mM for 3-MA, 10 µM for BBM and 100 nM for RAPA) was selected for subsequent experiments.

For integrin modulation, aggrecan (Sigma-Aldrich, A1960; 50 µg/mL) and manganese (5% solution prepared from manganese powder [Carl Roth, 1HX8.1] dissolved in HCl) were used. Treatments were applied either individually or in combination for 3 hours prior to analysis.

### Immunocytochemistry

Explants were fixed with 4% paraformaldehyde (PFA) for 15 minutes and permeabilized/blocking was performed using 1% goat serum and 2% Triton X-100 in PBS for 2 hours. Samples were then incubated overnight at 4 °C with the following primary antibodies: mouse monoclonal anti-βIII tubulin (Abcam, ab78078; 1:1200), rabbit polyclonal anti-LC3 (Cell Signaling Technology, 2775; 1:100), and rat monoclonal anti-LAMP1 (Abcam, ab25245; 1:100). The following day, explants were washed and incubated for 2 hours with secondary antibodies (1:400): goat anti-mouse Alexa Fluor 405, goat anti-rabbit Alexa Fluor 594, and goat anti-rat Alexa Fluor 488 (Thermo Fisher Scientific). Imaging was performed using an Andor Dragonfly 503 spinning disk confocal microscope equipped with a 63× objective, focusing on the distal region of the axon. Puncta quantification and colocalization analysis were carried out using the ComDet plugin in ImageJ (National Institutes of Health). Statistical analysis was conducted using GraphPad Prism 9. Data were tested for normality using the D’Agostino–Pearson test, and statistical significance was assessed using one-way ANOVA followed by Tukey’s post hoc test.

### Live imaging

Prior to live imaging, explants were incubated in Hibernate medium (Thermo Fisher Scientific, A1247501), supplemented as described above, either without or with Alexa Fluor® 647-conjugated anti-mouse/rat CD29 antibody (BioLegend, 102213; 1:100) for 20 minutes at room temperature to minimize integrin internalization. The anti-β1 integrin antibody used for live imaging exhibits integrin-blocking activity and modestly altered autophagic dynamics, consistent with the effects observed following aggrecan treatment (Fig. 6). As all experimental groups were imaged under identical antibody exposure and acquisition conditions, comparisons were performed only between axons exposed to the same labeling protocol. Following incubation, explants were washed several times with PBS. Explants were then either left intact or axotomized using a 23G syringe, generating a clear demarcation on the tissue surface. Subsequently, FluoroBrite™ DMEM-based medium (Thermo Fisher Scientific, A1896701), supplemented as described above, was added to reduce background during live imaging acquisition.

Live imaging was performed using an inverted spinning disk confocal microscope (Olympus SpinSR10) equipped with a 63× objective. Time-lapse acquisition was carried out at 1 frame per second for 2 minutes. DRG explants were imaged in a controlled chamber maintaining 5% CO₂, 37 °C, and appropriate humidity. For each experiment, live imaging was conducted in two configurations: (i) simultaneous green and red channels to assess autophagy dynamics (distinguishing AP and AL), and (ii) simultaneous red and far-red channels to evaluate integrin recycling (visualizing autophagic vesicles and integrins). Simultaneous acquisition of all three channels (green, red, and far-red) was avoided due to increased phototoxicity. Imaging was focused on the distal axon region (∼100 µm from the axon tip).

Data analysis was performed using ImageJ (National Institutes of Health). Kymographs were generated using the Multi Kymograph plugin, and vesicles were manually traced. The total number of vesicles observed over the 2-minute recording was quantified and normalized to the length of the analyzed axon segment (µm). Vesicles were classified based on their trajectory as anterograde (movement >5 µm/2 min toward the axon tip), retrograde (movement >5 µm/2 min toward the soma), or stationary (movement <5 µm/2 min).

For growth cone analysis, regions of interest (ROIs) were defined, and vesicles were tracked using the TrackMate plugin in ImageJ. The “maximum track displacement” parameter was used to quantify the maximum distance (µm) travelled by vesicles within the growth cone. Comparisons between pre- and post-injury growth cones were performed using an unpaired t-test.

Statistical analyses were conducted using GraphPad Prism 9. Data were tested for normality using the D’Agostino–Pearson test. Total vesicle number per axon was analyzed using one-way ANOVA followed by Tukey’s post hoc test. Vesicle proportion (AP:AL) and motility distributions (stationary, retrograde, anterograde) were calculated per axon as percentages of total vesicles and analyzed using two-way ANOVA with factors condition and movement type, followed by Tukey’s multiple comparisons test. A minimum of three independent experiments were performed for data acquisition.

### DRG explants neurite growth and regeneration assay

On day *in vitro* 3 (DIV3), explants were treated with autophagy-modulating compounds as described above. Axons were either left uncut or axotomized using a 23G syringe, generating a clear demarcation on the tissue surface. An initial image acquisition was performed using an Axio Observer D1 inverted phase-contrast microscope (Zeiss). Explants were then returned to the incubator, and a second image acquisition was performed 2 hours later. A minimum of three independent experiments per condition was conducted. Neurite growth (uncut) or regeneration (cut) was quantified using ImageJ (National Institutes of Health) defined as axonal elongation >5 µm between the two time points. Statistical analysis was performed using GraphPad Prism 9. Data were analyzed using one-way ANOVA followed by Tukey’s post hoc test.

## AUTHOR CONTRIBUTIONS

CRediT: Anda Cimpean: Conceptualization, Data curation, Formal analysis, Investigation, Methodology, Visualization, Writing - original draft, Writing - review & editing; Jessica Kwok: Supervision, Writing - review & editing; James W. Fawcett: Resources, Supervision, Writing - review & editing; Pavla Jendelova: Funding acquisition, Resources, Supervision, Writing - review & editing.

## DATA AVAILABILITY STATEMENT

The data that support the findings of this study are available at https://zenodo.org/records/21284658?preview=1&token=eyJhbGciOiJIUzUxMiJ9.eyJpZCI6I mMzYTRlN2YyLTRlOTYtNDNjMS04YmJkLTY5MTNjNTBlMzYxNCIsImRhdGEiOnt9L CJyYW5kb20iOiI5NTQxN2RlODc2MGUxNmYwMmEyZDUwOWY4NDhiNGQ1OCJ9.Q bBgYbwgbzBGTuvqPg4BD83JivMTxK9TrHv0Mo06uvDLW3fwEFlCkOH-tUPcdIJ2BfdboL7eZWlsdNCo9jBwIg

## ABBREVIATIONS

3-MA: 3-methyladenine
autolysosome: AL
autophagosome: AP
BBM: berbamine
CSPGs: chondroitin sulfate proteoglycans
DRG: dorsal root ganglia
ECM: extracellular matrix
FA: focal adhesion
Mn²+: manganese
RAPA: rapamycin

