## Supplementary material for "Intertwined Autophagy and Integrin Dynamics Shape Axon Growth and Regeneration": Table S1

**Supplementary table 1.** Statistics figure 2.

| **Total nr. LC3 vesicles** | **Mean 1** | **Mean 2** | **P Value** |
|---|---|---|---|
| **GROWING vs. STATIC** | 17.02 | 18.91 | 0.1537 |
| **Total nr. β1 integrin vesicles** | **Mean 1** | **Mean 2** | **P Value** |
| **GROWING vs. STATIC** | 29.48 | 30.77 | 0.6886 |
| **% coloc. LC3/β1 vesicles** | **Mean 1** | **Mean 2** | **P Value** |
| **GROWING vs. STATIC** | 19.04 | 22.11 | 0.4598 |

| **Proportion of LC3 vesicles** | | | |
|---|---|---|---|
| **GROWING vs. STATIC** | **Mean 1** | **Mean 2** | **P Value** |
| **AUTOPHAGOSOMES** | 37.86 | 65.1 | **<0.0001** |
| **AUTOLYSOSOMES** | 62.14 | 34.92 | **<0.0001** |

| **Autophagosomes movement** | | | |
|---|---|---|---|
| **GROWING vs. STATIC** | **Mean 1** | **Mean 2** | **P Value** |
| **ANTEROGRADE** | 5.536 | 4.21 | 0.9819 |
| **RETROGRADE** | 23.98 | 10.98 | **0.0034** |
| **STATIONARY** | 70.5 | 84.62 | **0.0013** |

| **Autolysosomes movement** | | | |
|---|---|---|---|
| **GROWING vs. STATIC** | **Mean 1** | **Mean 2** | **P Value** |
| **ANTEROGRADE** | 19.27 | 17.57 | 0.9813 |
| **RETROGRADE** | 31.02 | 31.37 | 0.9998 |
| **STATIONARY** | 49.7 | 52.79 | 0.8977 |

| **β1 vesicles movement** | | | |
|---|---|---|---|
| **GROWING vs. STATIC** | **Mean 1** | **Mean 2** | **P Value** |
| **ANTEROGRADE** | 5.758 | 3.258 | 0.5598 |
| **RETROGRADE** | 8.662 | 5.031 | 0.2416 |
| **STATIONARY** | 85.58 | 91.71 | **0.0127** |

| **Coloc. LC3/β1 vesicles movement** | | | |
|---|---|---|---|
| **GROWING vs. STATIC** | **Mean 1** | **Mean 2** | **P Value** |
| **ANTEROGRADE** | 29.71 | 1.42E-14 | **0.0302** |
| **RETROGRADE** | 14.48 | 45.42 | **0.018** |
| **STATIONARY** | 20.05 | 25.18 | 0.8744 |

**Supplementary table 2.** Statistics figure 3.

| **Total nr. LC3 vesicles** | **Mean 1** | **Mean 2** | **P Value** |
|---|---|---|---|
| BEFORE CUT vs. REG. CUT | 17.02 | 20.47 | **0.0281** |
| BEFORE CUT vs. STATIC CUT | 17.02 | 20.39 | **0.0396** |
| BEFORE CUT vs. RETR. CUT | 17.02 | 22.59 | **0.033** |
| REG. CUT vs. STATIC CUT | 20.47 | 20.39 | >0.9999 |
| REG. CUT vs. RETR. CUT | 20.47 | 22.59 | 0.7326 |
| STATIC CUT vs. RETR. CUT | 20.39 | 22.59 | 0.7132 |

| **Total nr. β1 integrin vesicles** | **Mean 1** | **Mean 2** | **P Value** |
|---|---|---|---|
| BEFORE CUT vs. REG. CUT | 29.48 | 40.98 | **0.0014** |
| BEFORE CUT vs. STATIC CUT | 29.48 | 38.1 | **0.0479** |
| BEFORE CUT vs. RETR. CUT | 29.48 | 41.25 | **0.04** |
| REG. CUT vs. STATIC CUT | 40.98 | 38.1 | 0.829 |
| REG. CUT vs. RETR. CUT | 40.98 | 41.25 | >0.9999 |
| STATIC CUT vs. RETR. CUT | 38.1 | 41.25 | 0.9034 |

| **% coloc. LC3/β1 vesicles** | **Mean 1** | **Mean 2** | **P Value** |
|---|---|---|---|
| BEFORE CUT vs. REG. CUT | 19.04 | 12.71 | 0.2111 |
| BEFORE CUT vs. STATIC CUT | 19.04 | 22.34 | 0.7684 |
| BEFORE CUT vs. RETR. CUT | 19.04 | 27.36 | 0.2596 |
| REG. CUT vs. STATIC CUT | 12.71 | 22.34 | **0.0384** |
| REG. CUT vs. RETR. CUT | 12.71 | 27.36 | **0.0106** |
| STATIC CUT vs. RETR. CUT | 22.34 | 27.36 | 0.715 |

| **Proportion of LC3 vesicles** | | | |
|---|---|---|---|
| **AUTOPHAGOSOMES** | **Mean 1** | **Mean 2** | **P Value** |
| BEFORE CUT vs. REG. CUT | 37.86 | 58.94 | **0.0003** |
| BEFORE CUT vs. STATIC CUT | 37.86 | 68.32 | **<0.0001** |
| BEFORE CUT vs. RETR. CUT | 37.86 | 72.08 | **0.0003** |
| REG. CUT vs. STATIC CUT | 58.94 | 68.32 | 0.2705 |
| REG. CUT vs. RETR. CUT | 58.94 | 72.08 | 0.3834 |
| STATIC CUT vs. RETR. CUT | 68.32 | 72.08 | 0.9692 |
| **AUTOLYSOSOMES** | **Mean 1** | **Mean 2** | **P Value** |
| BEFORE CUT vs. REG. CUT | 62.14 | 41.06 | **0.0003** |
| BEFORE CUT vs. STATIC CUT | 62.14 | 32.17 | **<0.0001** |
| BEFORE CUT vs. RETR. CUT | 62.14 | 27.92 | **0.0003** |
| REG. CUT vs. STATIC CUT | 41.06 | 32.17 | 0.317 |
| REG. CUT vs. RETR. CUT | 41.06 | 27.92 | 0.3834 |
| STATIC CUT vs. RETR. CUT | 32.17 | 27.92 | 0.9565 |

| Autophagosomes movement | | | |
| --- | --- | --- | --- |
| **ANTEROGRADE** | **Mean 1** | **Mean 2** | **P Value** |
| BEFORE CUT vs. REG. CUT | 5.536 | 4.01 | 0.9729 |
| BEFORE CUT vs. STATIC CUT | 5.536 | 3.394 | 0.9346 |
| BEFORE CUT vs. RETR. CUT | 5.536 | 10.82 | 0.7898 |
| REG. CUT vs. STATIC CUT | 4.01 | 3.394 | 0.9982 |
| REG. CUT vs. RETR. CUT | 4.01 | 10.82 | 0.6283 |
| STATIC CUT vs. RETR. CUT | 3.394 | 10.82 | 0.5677 |
| **RETROGRADE** | **Mean 1** | **Mean 2** | **P Value** |
| BEFORE CUT vs. REG. CUT | 23.98 | 7.768 | **<0.0001** |
| BEFORE CUT vs. STATIC CUT | 23.98 | 10.65 | **0.0015** |
| BEFORE CUT vs. RETR. CUT | 23.98 | 6.299 | **0.0109** |
| REG. CUT vs. STATIC CUT | 7.768 | 10.65 | 0.8551 |
| REG. CUT vs. RETR. CUT | 7.768 | 6.299 | 0.9939 |
| STATIC CUT vs. RETR. CUT | 10.65 | 6.299 | 0.8738 |
| **STATIONARY** | **Mean 1** | **Mean 2** | **P Value** |
| BEFORE CUT vs. REG. CUT | 70.5 | 88.28 | **<0.0001** |
| BEFORE CUT vs. STATIC CUT | 70.5 | 85.98 | **0.0001** |
| BEFORE CUT vs. RETR. CUT | 70.5 | 82.96 | 0.1286 |
| REG. CUT vs. STATIC CUT | 88.28 | 85.98 | 0.9198 |
| REG. CUT vs. RETR. CUT | 88.28 | 82.96 | 0.7858 |
| STATIC CUT vs. RETR. CUT | 85.98 | 82.96 | 0.9528 |

| Autolysosomes movement | | | |
| --- | --- | --- | --- |
| **ANTEROGRADE** | **Mean 1** | **Mean 2** | **P Value** |
| BEFORE CUT vs. REG. CUT | 19.27 | 22.85 | 0.861 |
| BEFORE CUT vs. STATIC CUT | 19.27 | 16.8 | 0.9574 |
| BEFORE CUT vs. RETR. CUT | 19.27 | 26.82 | 0.7474 |
| REG. CUT vs. STATIC CUT | 22.85 | 16.8 | 0.6174 |
| REG. CUT vs. RETR. CUT | 22.85 | 26.82 | 0.9537 |
| STATIC CUT vs. RETR. CUT | 16.8 | 26.82 | 0.5716 |
| **RETROGRADE** | **Mean 1** | **Mean 2** | **P Value** |
| BEFORE CUT vs. REG. CUT | 31.02 | 15.29 | **0.0034** |
| BEFORE CUT vs. STATIC CUT | 31.02 | 15.66 | **0.0094** |
| BEFORE CUT vs. RETR. CUT | 31.02 | 34.27 | 0.9729 |
| REG. CUT vs. STATIC CUT | 15.29 | 15.66 | 0.9998 |
| REG. CUT vs. RETR. CUT | 15.29 | 34.27 | 0.0611 |
| STATIC CUT vs. RETR. CUT | 15.66 | 34.27 | 0.0802 |
| **STATIONARY** | **Mean 1** | **Mean 2** | **P Value** |
| BEFORE CUT vs. REG. CUT | 49.7 | 61.9 | **0.0388** |
| BEFORE CUT vs. STATIC CUT | 49.7 | 67.54 | **0.0016** |
| BEFORE CUT vs. RETR. CUT | 49.7 | 38.91 | 0.4778 |
| REG. CUT vs. STATIC CUT | 61.9 | 67.54 | 0.6702 |
| REG. CUT vs. RETR. CUT | 61.9 | 38.91 | 0.0138 |
| STATIC CUT vs. RETR. CUT | 67.54 | 38.91 | 0.0015 |

| β1 vesicles movement | | | |
|---|---|---|---|
| **ANTEROGRADE** | **Mean 1** | **Mean 2** | **P Value** |
| BEFORE CUT vs. REG. CUT | 5.758 | 2.31 | 0.1301 |
| BEFORE CUT vs. STATIC CUT | 5.758 | 3.83 | 0.6642 |
| BEFORE CUT vs. RETR. CUT | 5.758 | 2.007 | 0.3519 |
| REG. CUT vs. STATIC CUT | 2.31 | 3.83 | 0.8187 |
| REG. CUT vs. RETR. CUT | 2.31 | 2.007 | 0.9992 |
| STATIC CUT vs. RETR. CUT | 3.83 | 2.007 | 0.8708 |
| **RETROGRADE** | **Mean 1** | **Mean 2** | **P Value** |
| BEFORE CUT vs. REG. CUT | 8.662 | 2.234 | **0.0003** |
| BEFORE CUT vs. STATIC CUT | 8.662 | 1.972 | **0.0005** |
| BEFORE CUT vs. RETR. CUT | 8.662 | 1.74 | **0.0134** |
| REG. CUT vs. STATIC CUT | 2.234 | 1.972 | 0.9988 |
| REG. CUT vs. RETR. CUT | 2.234 | 1.74 | 0.9966 |
| STATIC CUT vs. RETR. CUT | 1.972 | 1.74 | 0.9997 |
| **STATIONARY** | **Mean 1** | **Mean 2** | **P Value** |
| BEFORE CUT vs. REG. CUT | 85.58 | 95.46 | **<0.0001** |
| BEFORE CUT vs. STATIC CUT | 85.58 | 94.2 | **<0.0001** |
| BEFORE CUT vs. RETR. CUT | 85.58 | 96.25 | **<0.0001** |
| REG. CUT vs. STATIC CUT | 95.46 | 94.2 | 0.8879 |
| REG. CUT vs. RETR. CUT | 95.46 | 96.25 | 0.9859 |
| STATIC CUT vs. RETR. CUT | 94.2 | 96.25 | 0.8252 |

**Supplementary table 3.** Statistics figure 4.

| Total nr. LC3 vesicles | Mean 1 | Mean 2 | P Value |
|---|---|---|---|
| BEFORE CUT vs. REG | 17.02 | 15.38 | 0.5612 |
| BEFORE CUT vs. STATIC | 17.02 | 17.69 | 0.9599 |
| BEFORE CUT vs. RETR | 17.02 | 25.27 | **0.0009** |
| REG vs. STATIC | 15.38 | 17.69 | 0.2853 |
| REG vs. RETR | 15.38 | 25.27 | **<0.0001** |
| STATIC vs. RETR | 17.69 | 25.27 | **0.003** |

| Total nr. β1 integrin vesicles | Mean 1 | Mean 2 | P Value |
|---|---|---|---|
| BEFORE CUT vs. REG | 29.48 | 42.64 | **0.0006** |
| BEFORE CUT vs. STATIC | 29.48 | 40.04 | **0.0151** |
| BEFORE CUT vs. RETR | 29.48 | 38.58 | 0.266 |
| REG vs. STATIC | 42.64 | 40.04 | 0.8887 |
| REG vs. RETR | 42.64 | 38.58 | 0.8548 |
| STATIC vs. RETR | 40.04 | 38.58 | 0.9923 |

| % coloc. LC3/β1 vesicles | Mean 1 | Mean 2 | P Value |
|---|---|---|---|
| BEFORE CUT vs. REG | 19.04 | 10.95 | **0.0326** |
| BEFORE CUT vs. STATIC | 19.04 | 18.03 | 0.9848 |
| BEFORE CUT vs. RETR | 19.04 | 26.73 | 0.2164 |
| REG vs. STATIC | 10.95 | 18.03 | 0.1191 |
| REG vs. RETR | 10.95 | 26.73 | **0.0015** |
| STATIC vs. RETR | 18.03 | 26.73 | 0.1593 |

| Proportion of LC3 vesicles | | | |
|---|---|---|---|
| **AUTOPHAGOSOMES** | **Mean 1** | **Mean 2** | **P Value** |
| BEFORE CUT vs. REG | 37.86 | 46.34 | 0.3199 |
| BEFORE CUT vs. STATIC | 37.86 | 58.27 | **0.0006** |
| BEFORE CUT vs. RETR | 37.86 | 42.44 | 0.9465 |
| REG vs. STATIC | 46.34 | 58.27 | 0.0753 |
| REG vs. RETR | 46.34 | 42.44 | 0.9639 |
| STATIC vs. RETR | 58.27 | 42.44 | 0.2276 |
| **AUTOLYSOSOMES** | **Mean 1** | **Mean 2** | **P Value** |
| BEFORE CUT vs. REG | 62.14 | 53.66 | 0.3199 |
| BEFORE CUT vs. STATIC | 62.14 | 41.75 | **0.0006** |
| BEFORE CUT vs. RETR | 62.14 | 57.64 | 0.9492 |
| REG vs. STATIC | 53.66 | 41.75 | 0.0761 |
| REG vs. RETR | 53.66 | 57.64 | 0.9617 |
| STATIC vs. RETR | 41.75 | 57.64 | 0.2244 |

| Autophagosomes movement | | | |
|---|---|---|---|
| **ANTEROGRADE** | **Mean 1** | **Mean 2** | **P Value** |
| BEFORE CUT vs. REG | 5.536 | 1.614 | 0.6027 |
| BEFORE CUT vs. STATIC | 5.536 | 1.998 | 0.7217 |
| BEFORE CUT vs. RETR | 5.536 | 1.364 | 0.877 |
| REG vs. STATIC | 1.614 | 1.998 | 0.9994 |
| REG vs. RETR | 1.614 | 1.364 | >0.9999 |
| STATIC vs. RETR | 1.998 | 1.364 | 0.9995 |
| **RETROGRADE** | **Mean 1** | **Mean 2** | **P Value** |
| BEFORE CUT vs. REG | 23.98 | 7.952 | **<0.0001** |
| BEFORE CUT vs. STATIC | 23.98 | 5.624 | **<0.0001** |
| BEFORE CUT vs. RETR | 23.98 | 22.75 | 0.9962 |
| REG vs. STATIC | 7.952 | 5.624 | 0.8864 |
| REG vs. RETR | 7.952 | 22.75 | **0.0351** |
| STATIC vs. RETR | 5.624 | 22.75 | **0.0123** |
| **STATIONARY** | **Mean 1** | **Mean 2** | **P Value** |
| BEFORE CUT vs. REG | 70.5 | 90.42 | **<0.0001** |
| BEFORE CUT vs. STATIC | 70.5 | 92.4 | **<0.0001** |
| BEFORE CUT vs. RETR | 70.5 | 75.88 | 0.7689 |
| REG vs. STATIC | 90.42 | 92.4 | 0.9262 |
| REG vs. RETR | 90.42 | 75.88 | **0.0401** |
| STATIC vs. RETR | 92.4 | 75.88 | **0.0172** |

| Autolysosomes movement | | | |
|---|---|---|---|
| **ANTEROGRADE** | **Mean 1** | **Mean 2** | **P Value** |
| BEFORE CUT vs. REG | 19.29 | 15.33 | 0.7425 |
| BEFORE CUT vs. STATIC | 19.29 | 12.58 | 0.3527 |
| BEFORE CUT vs. RETR | 19.29 | 16.8 | 0.9824 |
| REG vs. STATIC | 15.33 | 12.58 | 0.9037 |
| REG vs. RETR | 15.33 | 16.8 | 0.9962 |
| STATIC vs. RETR | 12.58 | 16.8 | 0.9236 |
| **RETROGRADE** | **Mean 1** | **Mean 2** | **P Value** |
| BEFORE CUT vs. REG | 31.02 | 17.46 | **0.0032** |
| BEFORE CUT vs. STATIC | 31.02 | 9.291 | **<0.0001** |
| BEFORE CUT vs. RETR | 31.02 | 23.66 | 0.6877 |
| REG vs. STATIC | 17.46 | 9.291 | 0.1807 |
| REG vs. RETR | 17.46 | 23.66 | 0.7883 |
| STATIC vs. RETR | 9.291 | 23.66 | 0.1459 |
| **STATIONARY** | **Mean 1** | **Mean 2** | **P Value** |
| BEFORE CUT vs. REG | 49.73 | 67.19 | **<0.0001** |
| BEFORE CUT vs. STATIC | 49.73 | 78.13 | **<0.0001** |
| BEFORE CUT vs. RETR | 49.73 | 59.54 | 0.4569 |
| REG vs. STATIC | 67.19 | 78.13 | **0.0349** |
| REG vs. RETR | 67.19 | 59.54 | 0.659 |
| STATIC vs. RETR | 78.13 | 59.54 | **0.031** |

| β1 vesicles movement | | | |
|---|---|---|---|
| **ANTEROGRADE** | **Mean 1** | **Mean 2** | **P Value** |
| BEFORE CUT vs. REG | 5.758 | 4.143 | 0.7367 |
| BEFORE CUT vs. STATIC | 5.758 | 3.299 | 0.4614 |
| BEFORE CUT vs. RETR | 5.758 | 4.566 | 0.9509 |
| REG vs. STATIC | 4.143 | 3.299 | 0.9632 |
| REG vs. RETR | 4.143 | 4.566 | 0.9978 |
| STATIC vs. RETR | 3.299 | 4.566 | 0.9501 |
| **RETROGRADE** | **Mean 1** | **Mean 2** | **P Value** |
| BEFORE CUT vs. REG | 8.662 | 5.203 | 0.1283 |
| BEFORE CUT vs. STATIC | 8.662 | 2.998 | **0.0047** |
| BEFORE CUT vs. RETR | 8.662 | 6.18 | 0.6832 |
| REG vs. STATIC | 5.203 | 2.998 | 0.5913 |
| REG vs. RETR | 5.203 | 6.18 | 0.9739 |
| STATIC vs. RETR | 2.998 | 6.18 | 0.5334 |
| **STATIONARY** | **Mean 1** | **Mean 2** | **P Value** |
| BEFORE CUT vs. REG | 85.58 | 90.65 | **0.0079** |
| BEFORE CUT vs. STATIC | 85.58 | 93.7 | **<0.0001** |
| BEFORE CUT vs. RETR | 85.58 | 89.84 | 0.2274 |
| REG vs. STATIC | 90.65 | 93.7 | 0.306 |
| REG vs. RETR | 90.65 | 89.84 | 0.9846 |
| STATIC vs. RETR | 93.7 | 89.84 | 0.3603 |

| Coloc. LC3/β1 vesicles movement | | | |
|---|---|---|---|
| **ANTEROGRADE** | **Mean 1** | **Mean 2** | **P Value** |
| BEFORE CUT vs. REG | 29.71 | 18.6 | 0.5952 |
| BEFORE CUT vs. STATIC | 29.71 | 26.19 | 0.9837 |
| BEFORE CUT vs. RETR | 29.71 | 20 | 0.9021 |
| REG vs. STATIC | 18.6 | 26.19 | 0.8756 |
| REG vs. RETR | 18.6 | 20 | 0.9997 |
| STATIC vs. RETR | 26.19 | 20 | 0.9759 |
| **RETROGRADE** | **Mean 1** | **Mean 2** | **P Value** |
| BEFORE CUT vs. REG | 14.48 | 37.1 | **0.029** |
| BEFORE CUT vs. STATIC | 14.48 | 27.27 | 0.5892 |
| BEFORE CUT vs. RETR | 14.48 | 28.57 | 0.6481 |
| REG vs. STATIC | 37.1 | 27.27 | 0.7896 |
| REG vs. RETR | 37.1 | 28.57 | 0.9025 |
| STATIC vs. RETR | 27.27 | 28.57 | 0.9997 |
| **STATIONARY** | **Mean 1** | **Mean 2** | **P Value** |
| BEFORE CUT vs. REG | 20.05 | 13.77 | 0.805 |
| BEFORE CUT vs. STATIC | 20.05 | 17.67 | 0.9886 |
| BEFORE CUT vs. RETR | 20.05 | 24.4 | 0.9711 |
| REG vs. STATIC | 13.77 | 17.67 | 0.9575 |
| REG vs. RETR | 13.77 | 24.4 | 0.7163 |
| STATIC vs. RETR | 17.67 | 24.4 | 0.916 |

**Supplementary table 4.** Statistics figure 5.

| MAX. TRACK DISPLACEMENT LC3+ VESICLES | Mean 1 | Mean 2 | P Value |
|---|---|---|---|
| GC BEFORE CUT vs. GC AFTER CUT | 0.5739 | 0.4422 | **<0.0001** |

| MAX. TRACK DISPLACEMENT β1+ VESICLES | Mean 1 | Mean 2 | P Value |
|---|---|---|---|
| GC BEFORE CUT vs. GC AFTER CUT | 0.2914 | 0.1939 | **0.0409** |

**Supplementary table 5.** Statistics figure 6.

| Total nr. LC3 vesicles | Mean 1 | Mean 2 | P Value |
|---|---|---|---|
| CONTROL vs. Aggrecan | 16.12 | 14.03 | 0.5163 |
| CONTROL vs. Mn | 16.12 | 17.33 | 0.8837 |
| CONTROL vs. Aggrencan+Mn | 16.12 | 14.38 | 0.6374 |
| Aggrecan vs. Mn | 14.03 | 17.33 | 0.1608 |
| Aggrecan vs. Aggrencan+Mn | 14.03 | 14.38 | 0.9938 |
| Mn vs. Aggrencan+Mn | 17.33 | 14.38 | 0.2196 |

| Proportion of LC3 vesicles | | | |
|---|---|---|---|
| **AUTOPHAGOSOMES** | **Mean 1** | **Mean 2** | **P Value** |
| CONTROL vs. Aggrecan | 34.88 | 56.55 | **0.018** |
| CONTROL vs. Mn | 34.88 | 47.31 | 0.4181 |
| CONTROL vs. Aggrecan+Mn | 34.88 | 30.08 | 0.9107 |
| Aggrecan vs. Mn | 56.55 | 47.31 | 0.6007 |
| Aggrecan vs. Aggrecan+Mn | 56.55 | 30.08 | **0.0004** |
| Mn vs. Aggrecan+Mn | 47.31 | 30.08 | 0.0943 |
| **AUTOLYSOSOMES** | **Mean 1** | **Mean 2** | **P Value** |
| CONTROL vs. Aggrecan | 65.18 | 43.45 | **0.0176** |
| CONTROL vs. Mn | 65.18 | 52.75 | 0.4184 |
| CONTROL vs. Aggrecan+Mn | 65.18 | 69.92 | 0.9136 |
| Aggrecan vs. Mn | 43.45 | 52.75 | 0.5954 |
| Aggrecan vs. Aggrecan+Mn | 43.45 | 69.92 | **0.0004** |
| Mn vs. Aggrecan+Mn | 52.75 | 69.92 | 0.0962 |

| Autophagosomes movement | | | |
|---|---|---|---|
| **ANTEROGRADE** | **Mean 1** | **Mean 2** | **P Value** |
| CONTROL vs. Aggrecan | 4.533 | 9.781 | 0.8514 |
| CONTROL vs. Mn | 4.533 | 8.533 | 0.9304 |
| CONTROL vs. Aggrecan+Mn | 4.533 | 6.522 | 0.9875 |
| Aggrecan vs. Mn | 9.781 | 8.533 | 0.9975 |
| Aggrecan vs. Aggrecan+Mn | 9.781 | 6.522 | 0.9457 |
| Mn vs. Aggrecan+Mn | 8.533 | 6.522 | 0.987 |
| **RETROGRADE** | **Mean 1** | **Mean 2** | **P Value** |
| CONTROL vs. Aggrecan | 20.13 | 18.32 | 0.9924 |
| CONTROL vs. Mn | 20.13 | 36.47 | 0.0679 |
| CONTROL vs. Aggrecan+Mn | 20.13 | 8.739 | 0.2332 |
| Aggrecan vs. Mn | 18.32 | 36.47 | **0.0295** |
| Aggrecan vs. Aggrecan+Mn | 18.32 | 8.739 | 0.3667 |
| Mn vs. Aggrecan+Mn | 36.47 | 8.739 | **<0.0001** |
| **STATIONARY** | **Mean 1** | **Mean 2** | **P Value** |
| CONTROL vs. Aggrecan | 75.27 | 71.9 | 0.9548 |
| CONTROL vs. Mn | 75.27 | 55.07 | **0.0136** |
| CONTROL vs. Aggrecan+Mn | 75.27 | 84.78 | 0.3905 |
| Aggrecan vs. Mn | 71.9 | 55.07 | 0.0507 |
| Aggrecan vs. Aggrecan+Mn | 71.9 | 84.78 | 0.1308 |
| Mn vs. Aggrecan+Mn | 55.07 | 84.78 | **<0.0001** |

| Autolysosomes movement | | | |
|---|---|---|---|
| **ANTEROGRADE** | **Mean 1** | **Mean 2** | **P Value** |
| CONTROL vs. Aggrecan | 15.76 | 19.84 | 0.9508 |
| CONTROL vs. Mn | 15.76 | 31.38 | 0.2071 |
| CONTROL vs. Aggrecan+Mn | 15.76 | 26.69 | 0.4208 |
| Aggrecan vs. Mn | 19.84 | 31.38 | 0.4482 |
| Aggrecan vs. Aggrecan+Mn | 19.84 | 26.69 | 0.7543 |
| Mn vs. Aggrecan+Mn | 31.38 | 26.69 | 0.9176 |
| **RETROGRADE** | **Mean 1** | **Mean 2** | **P Value** |
| CONTROL vs. Aggrecan | 33.12 | 16.16 | 0.1211 |
| CONTROL vs. Mn | 33.12 | 26.25 | 0.8246 |
| CONTROL vs. Aggrecan+Mn | 33.12 | 26.5 | 0.7904 |
| Aggrecan vs. Mn | 16.16 | 26.25 | 0.5641 |
| Aggrecan vs. Aggrecan+Mn | 16.16 | 26.5 | 0.4409 |
| Mn vs. Aggrecan+Mn | 26.25 | 26.5 | >0.9999 |
| **STATIONARY** | **Mean 1** | **Mean 2** | **P Value** |
| CONTROL vs. Aggrecan | 51.06 | 63.95 | 0.333 |
| CONTROL vs. Mn | 51.06 | 42.5 | 0.7059 |
| CONTROL vs. Aggrecan+Mn | 51.06 | 46.77 | 0.9317 |
| Aggrecan vs. Mn | 63.95 | 42.5 | **0.0314** |
| Aggrecan vs. Aggrecan+Mn | 63.95 | 46.77 | 0.0647 |
| Mn vs. Aggrecan+Mn | 42.5 | 46.77 | 0.9359 |

**Supplementary table 6.** Statistics figure 7.

| **Total nr. LC3 vesicles** | **Mean 1** | **Mean 2** | **P Value** |
|---|---|---|---|
| CONTROL vs. 3MA 1mM | 31.77 | 22.68 | **0.0021** |
| CONTROL vs. 3MA 2.5mM | 31.77 | 14.02 | **<0.0001** |
| CONTROL vs. BBM 1uM | 31.77 | 31.02 | 0.9816 |
| CONTROL vs. BBM 10uM | 31.77 | 39.37 | **0.0293** |
| CONTROL vs. RAPA 1nM | 31.77 | 38.66 | **0.0488** |
| CONTROL vs. RAPA 100nM | 31.77 | 48.29 | **0.0012** |

| Proportion of LC3 vesicles | | | |
|---|---|---|---|
| **AUTOPHAGOSOMES** | **Mean 1** | **Mean 2** | **P Value** |
| CONTROL vs. 3MA 1mM | 45.88 | 34.99 | 0.149 |
| CONTROL vs. 3MA 2.5mM | 45.88 | 51.93 | 0.3933 |
| CONTROL vs. BBM 1uM | 45.88 | 37.04 | 0.2353 |
| CONTROL vs. BBM 10uM | 45.88 | 64.09 | **<0.0001** |
| CONTROL vs. RAPA 1nM | 45.88 | 40.8 | 0.3629 |
| CONTROL vs. RAPA 100nM | 45.88 | 25.9 | **0.0023** |
| **AUTOLYSOSOMES** | **Mean 1** | **Mean 2** | **P Value** |
| CONTROL vs. 3MA 1mM | 54.12 | 65.01 | 0.1492 |
| CONTROL vs. 3MA 2.5mM | 54.12 | 48.08 | 0.3936 |
| CONTROL vs. BBM 1uM | 54.12 | 62.96 | 0.2354 |
| CONTROL vs. BBM 10uM | 54.12 | 35.91 | **<0.0001** |
| CONTROL vs. RAPA 1nM | 54.12 | 59.2 | 0.3622 |
| CONTROL vs. RAPA 100nM | 54.12 | 74.1 | **0.0023** |

**Supplementary table 7.** Statistics figure 8.

| **Total nr. LC3 vesicles** | **Mean 1** | **Mean 2** | **P Value** |
|---|---|---|---|
| CONTROL vs. 3MA 2.5mM | 13.68 | 8.507 | **0.0209** |
| CONTROL vs. BBM 10uM | 13.68 | 11.9 | 0.555 |
| CONTROL vs. RAPA 100nM | 13.68 | 23.98 | **<0.0001** |

| **Proportion of LC3 vesicles** | | | |
|---|---|---|---|
| **AUTOPHAGOSOMES** | **Mean 1** | **Mean 2** | **P Value** |
| CONTROL vs. 3MA 2.5mM | 37.85 | 31.17 | 0.8205 |
| CONTROL vs. BBM 10uM | 37.85 | 85.29 | **<0.0001** |
| CONTROL vs. RAPA 100nM | 37.85 | 19.79 | 0.1151 |
| **AUTOLYSOSOMES** | **Mean 1** | **Mean 2** | **P Value** |
| CONTROL vs. 3MA 2.5mM | 62.13 | 68.83 | 0.8191 |
| CONTROL vs. BBM 10uM | 62.13 | 14.71 | **<0.0001** |
| CONTROL vs. RAPA 100nM | 62.13 | 80.27 | 0.1127 |

**Supplementary table 8.** Statistics figure 9.

| **% GROWING AXONS** | **Mean 1** | **Mean 2** | **P Value** |
|---|---|---|---|
| CONTROL vs. 3MA | 82.18 | 12.51 | **<0.0001** |
| CONTROL vs. BBM | 82.18 | 54.32 | **0.0029** |
| CONTROL vs. RAPA | 82.18 | 93.63 | 0.3048 |

| **GROWTH LENGTH** | **Mean 1** | **Mean 2** | **P Value** |
|---|---|---|---|
| CONTROL vs. 3MA | 30.17 | 13.4 | **<0.0001** |
| CONTROL vs. BBM | 30.17 | 27.24 | 0.5948 |
| CONTROL vs. RAPA | 30.17 | 48.96 | **<0.0001** |

| **% REGENERATING AXONS** | **Mean 1** | **Mean 2** | **P Value** |
|---|---|---|---|
| CONTROL vs. 3MA | 36.5 | 7.667 | **<0.0001** |
| CONTROL vs. BBM | 36.5 | 15.2 | **0.0024** |
| CONTROL vs. RAPA | 36.5 | 48.9 | **0.0315** |

| **REGENERATION LENGTH** | **Mean 1** | **Mean 2** | **P Value** |
|---|---|---|---|
| CONTROL vs. 3MA | 34.36 | 13 | **0.0056** |
| CONTROL vs. BBM | 34.36 | 40.5 | 0.7159 |
| CONTROL vs. RAPA | 34.36 | 35.7 | 0.9889 |

**Supplementary table 9.** Statistics figure 10.

| **Total nr. LC3 vesicles** | **Mean 1** | **Mean 2** | **P Value** |
|---|---|---|---|
| CONTROL vs. RAPA | 16.27 | 20.12 | **0.0347** |

| **Total nr. β1 integrin vesicles** | **Mean 1** | **Mean 2** | **P Value** |
|---|---|---|---|
| CONTROL vs. RAPA | 29.3 | 36.04 | **0.0186** |

| **% coloc. LC3/β1 vesicles** | **Mean 1** | **Mean 2** | **P Value** |
|---|---|---|---|
| CONTROL vs. RAPA | 19.04 | 12.41 | **0.0192** |

| **β1 vesicles movement** | | | |
|---|---|---|---|
| CONTROL vs. RAPA | **Mean 1** | **Mean 2** | **P Value** |
| ANTEROGRADE | 5.758 | 3.749 | 0.596 |
| RETROGRADE | 8.662 | 5.585 | 0.2356 |
| STATIONARY | 85.58 | 90.67 | **0.0142** |

| **Coloc. LC3/β1 vesicles movement** | | | |
|---|---|---|---|
| CONTROL vs. RAPA | **Mean 1** | **Mean 2** | **P Value** |
| ANTEROGRADE | 29.71 | 29.17 | >0.9999 |
| RETROGRADE | 17.33 | 51.75 | **0.0004** |
| STATIONARY | 20.05 | 11.05 | 0.5814 |
